# Pathogenic human huntingtin expression causes prolific intramuscular aggregation, leading to nuclear, metabolic, and physiological dysregulation in striated muscle

**DOI:** 10.64898/2026.04.20.719674

**Authors:** Tadros A. Hana, Kiel G. Ormerod

**Affiliations:** University of Toledo, Neuroscience and Psychiatry Department, Toledo, OH, U.S.A

**Keywords:** Huntington’s Disease, Drosophila, skeletal muscle, metabolism

## Abstract

Huntington’s disease is caused by expansion of a CAG repeat in the human HTT gene, producing a mutant huntingtin protein that misfolds and forms intracellular aggregates. Although Huntington’s disease is primarily characterized as a neurodegenerative disorder, mutant huntingtin is ubiquitously expressed, and peripheral tissues such as skeletal muscle exhibit pathological abnormalities. To define the muscle-intrinsic consequences of pathogenic huntingtin expression, we expressed caspase-6 truncated pathogenic human huntingtin in body wall muscle of *Drosophila melanogaster* larvae and performed quantitative structural and functional analyses. Aggregate analysis revealed that fluorescence intensity increased with aggregate size while aggregate morphology became more irregular. Delaying transgene expression until later stages of larval development dramatically reduced aggregate number, demonstrating a strong temporal dependence of aggregate formation. Myonuclei were enlarged, misshapen, and exhibited significantly reduced fluorescence intensity, consistent with altered chromatin organization. Notably, huntingtin aggregates were observed within the nucleus, indicating that nuclear proteostasis is directly perturbed by pathogenic huntingtin in muscle cells. Despite these intracellular defects, muscle fiber shape and sarcomere organization were preserved, suggesting that contractile apparatus assembly is not overtly disrupted. In contrast, mitochondrial organization was severely affected, with extensive mitochondrial aggregation throughout muscle fibers, consistent with altered organelle homeostasis. Functional analyses demonstrated that pathogenic huntingtin expression significantly impaired neuromuscular performance. Larvae exhibited reduced excitatory junctional potentials and diminished muscle contractile force, indicating compromised synaptic transmission and muscle function. Together, these findings demonstrate that pathogenic human huntingtin expression in skeletal muscle is sufficient to drive widespread protein aggregation, nuclear and mitochondrial abnormalities, and functional deficits despite the absence of overt structural changes. Our results highlight the importance of muscle-intrinsic pathogenic mechanisms and provide a quantitative framework for understanding how mutant huntingtin disrupts cellular organization and physiology outside the nervous system.

## Introduction

Huntington’s disease (HD) is an autosomal dominant neurodegenerative disorder caused by expansion of a CAG trinucleotide repeat in exon 1 of the HTT gene, resulting in an elongated glutamine tract within the N-terminus of the huntingtin protein^1^. HD has been assigned to the class of human polyglutamine or CAG-repeat diseases, all of which are currently untreatable^2^. Historically, HD has been characterized as a disorder of the CNS, driven by progressive degeneration of striatal and cortical neurons, however, a growing body of evidence demonstrates that HD is a multi-system disease with substantial peripheral pathology^3,4^. Some of the most prominent non-neurological manifestations of HD are progressive weight loss, skeletal muscle atrophy, metabolic dysfunction, and impaired motor performance, features which cannot be fully explained solely by neuronal degeneration^5–10^.

Pathogenic huntingtin (P-htt) is ubiquitously expressed, including in skeletal muscle, where it has been detected in both patient biopsies and animal models of HD^9,10^. Skeletal muscle abnormalities in HD include fiber-type switching, mitochondrial dysfunction, reduced oxidative capacity, impaired contractile force, and altered neuromuscular junction integrity^6,10–14^. Although some of these changes may arise secondary to denervation or reduced physical activity, increasing evidence supports a cell-autonomous contribution of P-htt within muscle tissue itself^3,4,9,15^. Indeed, muscle-specific expression of P-htt in transgenic models recapitulates aspects of HD-associated muscle pathology, whereas peripheral silencing of *HTT* can ameliorate metabolic and functional deficits, further implicating direct effects of P-htt in muscle^16,17^.

At the molecular level, pathogenic huntingtin has been shown to disrupt molecular transport, transcriptional regulation, protein homeostasis, mitochondrial dynamics, and calcium handling, processes which are all essential for skeletal muscle integrity and performance^18–25^. Given the dominant energetic demand and structural specialization of myofibers, muscle tissue may be particularly susceptible to disturbances in proteostasis and bioenergetic balance. Moreover, skeletal muscle functions as a key regulator of systemic metabolism, suggesting that muscle-specific pathology may contribute to whole-body metabolic alterations observed in HD patients^26,27^. Despite increasing awareness of peripheral involvement in HD, the mechanisms by which pathogenic huntingtin drives muscle dysfunction remain incompletely understood. Elucidating these mechanisms is critical not only for understanding disease pathogenesis but also for developing comprehensive therapeutic strategies that address both central and peripheral manifestations. In this study, we investigate the effects of pathogenic huntingtin expression in skeletal muscle, focusing on its impact on muscle structure, function, and cellular homeostasis. By defining the muscle-intrinsic consequences of P-htt expression, we aim to clarify the contribution of peripheral pathology to overall disease progression in Huntington’s disease.

## Materials and Methods

### Husbandry

*Drosophila melanogaster* were reared on standard media (g/L: sucrose: 11, corn meal: 52, yeast: 27.2, agar: 7.2, 2.38: 4-hydroxybenzoate) at 22 °C at constant humidity on a 12:12 light: dark cycle. Flies of both sexes were used for all experiments. A list of all fly stocks used in the study is found in table 1. UAS-htt-Q15, UAS-htt-Q138, and UAS-myr-GCaMP8s were generously provided by Troy Littleton, Massachusetts Institute of Technology. All other fly lines were obtained from Bloomington Drosophila Stock Center (BDSC) or created in our laboratory.

**Table 1.** Fly lines, genotypes, and abbreviation used in this study.

| Full genotype | Abbreviation | BDSC Stock # | Reference |
| --- | --- | --- | --- |
| UAS- HttQ15-mRFP/+ | NP-htt | - | 18,19 |
| UAS- HttQ138-mRFP/+ | P-htt | - | 18,19 |
| G7-Gal4/+ ; UAS- HttQ15-mRFP/+ | G7 > NP-htt | - | - |
| G7-Gal4/+ ; UAS- HttQ138-mRFP/+ | G7 > P-htt | - | - |
| Elav-Gal4 |  | 458 | 28 |
| Mef2-Gal4/+ ; UAS- HttQ15-mRFP/+ | Mef2 > NP-htt | - | - |
| Mef2-Gal4/+ ; UAS- HttQ138-mRFP/+ | Mef2 > P-htt | - | - |
| Mef2-Gal4 | Mef2 | - | 29 |
| G7-Gal4 | G7 | - | 30 |
| UAS-myr-GCaMP-8s | GCaMP | - | 31 |

### Larval huntingtin aggregation imaging

Third instar larvae expressing either UAS-htt-Q15 or UAS-htt-Q138 via muscle drivers Mef2-Gal4, G7-Gal4, and 5053A-Gal4 were isolated from fly vials, and dissected in hemolymph-like (HL3.0) calcium-free saline (in mM): NaCl: 70; KCl: 5; MgCl_2_: 4; NaHCO_3_: 10; Trehalose: 5; Sucrose: 115; HEPES: 5. pH was adjusted to 7.18. All reagents and chemicals were obtained from Research Products International unless otherwise specified. Following dissection, larvae were fixed in 4% paraformaldehyde (TissuePro) for five minutes followed by a wash in phosphate-buffered saline (PBS), with 0.05% Triton X-100 (PBS-T), and blocked with 5% bovine serine albumin (BSA). Larvae were mounted in mounting media with DAPI (abcam, ab104139). All images were taken with an Olympus BX51W1, equipped with a Lumencor Fluorescence Light Engine, Hammamatsu Orca-fusion camera, and processed using Olympus Cellsens software. Z-stacks were acquired using an objective nanopositioner (Nano-F200S). All images were taken with an Olympus UPLFLN 60x oil-immersion objective. *Aggregate analysis*: 20-60 images were acquired per muscle at 0.4-0.6 micrometer steps to generate Z-stacks. Olympus CellSens was used to compress those Z-stacks into maximum Z-projections. Subsequently, these images were cropped to include only the muscle fiber analyzed using a brightfield overlay. The cropped images were processed using CellSens 2D deconvolution filter set to 20%. Aggregates were detected using the manual threshold fluorescence feature, and the dimmest detectable signal was used to set the minimum threshold value. All regions of interest (ROIs) smaller than 0.2 microns^2^ were removed as this was the smallest area of reproducible detection by the software. For each aggregate, the following metrics were generated: number, size (area), gray scale (fluorescence intensity), and shape factor (formula: 4π x area/perimeter^2^). Raw data were generated and output into comma-separated values (CSV) files, compiled and analyzed using Microsoft Excel. Figures were generated using GraphPad Prism.

### Immunohistochemistry

Wandering third instar larvae were dissected in modified, calcium-free HL3.0 saline. The larvae were then fixed for 30 minutes in 4 % paraformaldehyde and subsequently washed in PBS-T, and blocked with 5 % BSA. Larvae were then incubated with primary antibodies in PBST at 4°C overnight and washed three times for 15 min in PBS-T. Larvae were then incubated for 2 hours at room temperature in secondary antibodies, followed by three 15 min washes in PBS-T. All preparations were mounted in a DAPI containing mounting media (ab104139). Antibodies used for this study include the following: DyLight 649 conjugated anti-horseradish peroxidase (HRP), 1:750 (catalog #123–605–021, Jackson ImmunoResearch), filamentous actin probe, phalloidin conjugated to Alexa Fluor^™^ 488 (A12379, Thermo, 1:3000), Mitochondrial marker anti-ATP5A (Clone 15H4C4, abcam, 1:500), 4F3 anti-discs-large (AB 528203, DHSB, 1:500), goat anti-mouse Alexa Fluor 488 (ThermoFisher Scientific, catalog # A11001, 1:500). Fixed and mounted animals were imaged on an Olympus BX51W1 microscope using either a 20x Zeiss APOCHROMAT 0.75 or a UPLFLN 60x oil-immersion objective. Images were acquired with a Hammamatsu Orca-fusion camera, and processed using Olympus CellSens software. **Muscle structure:** Length and width of individual MF 4 muscles were quantified using the straight-line length feature in CellSens. Muscle length was measured along the midline, starting and ending at the insertion points of each muscle fiber. Width was measured across the midline equidistant from the insertion points. Muscle area was calculated as the product of length multiplied by width. Only muscle fibers from abdominal segments A3-A5 were measured. Sarcomere and I-band length were measured according to our previously established protocol, using the line profile tool on CellSens from 60x phalloidin stained MF4s^32^. Raw data from CellSens were exported as CSV files, compiled in excel, analyzed and graphed in GraphPad Prism**. Nuclear morphology and intranuclear aggregate accumulation:** Nuclei were manually selected from maximum Z-stack projection overlays of DAPI and anti-actin using the region of interest (ROI) measuring tool in CellSens and the following metrics were generated: number, size (area), and shape factor. Additionally, maximum Z-stack projection overlays of DAPI and RFP-positive huntingtin punctae were created, and each individual nucleus in MF 4 was cropped into individual images and assessed for intranuclear aggregates. If a nucleus was positive for htt protein aggregation the manual fluorescence threshold feature was used to detect the following values: number, size (area), gray scale (fluorescence intensity), and shape factor. Additionally, total aggregate load was calculated as the sum area of all RFP-positive aggregates inside the perimeter of htt positive nuclei divided by the DAPI area, then expressed as a percentage. Raw data generated in CellSens were output as CSV files, compiled and analyzed using Excel, and figures were created using GraphPad Prism. **Periodic Acid Stain (PAS):** Stain was conducted as previously outlined in Banerjee et al, (2021). Briefly, larvae were dissected in HL3.1, fixed in 4% PFA in PBS for 20 min at room temperature (RT). Tissue was washed PBS with 1% BSA, then incubated in periodic acid solution (1 g/dl) for 5 min at RT, then washed in 1% BSA in PBS with Schiff’s reagent for 15 at RT, followed by a 10 min wash in PBS with 1% BSA^33^. Tissue was then mounted in glycerol, and imaged using light microscopy at 20 and 60 x magnification using an Olympus BX51W1 microscope and imaged using a Hammamatsu Orca-fusion camera. **Oil-red-O stain:** Larvae were dissected in HL3.1, fixed in 4% PFA in PBS for 10 min, then washed in PBS with 1% BSA three times for 10 min each wash. Tissue was then incubated for 25 min in Oil Red O stain (6 mL of 0.1% Oil red O in isopropanol + 4 mL of dH_2_O, solution filtered through 0.45 micron filter syringe). Tissue was then rinsed three times for 10 minutes in PBS, mounted on microscope slides using glycerol, and imaged at 20 and 60 x magnification on our Olympus scope with Hammamatsu camera.

### Spontaneous release calcium imaging (GCaMP)

Wandering third instar larvae expressing postsynaptic (Mef2-Gal4) UAS-myr-GCaMP7 were dissected in calcium-free HL3.1 saline at room temperature. The preparation was washed several times with HL3.1 saline containing 1.3mM calcium, and allowed to rest for 5 minutes. NMJs from MF 4 from AS3-5 were imaged continuously for 1 minute at 10 frames per second (fps, 600 fames total) using an Olympus microscope equipped with a Lumencor Fluorescence light engine, hommamatsu Orca-fusion camera, using a 60x water immersion objective. Images were acquired using Olympus CellSens software, then raw images were exported as tiff files. Spontaneous release events were analyzed using Volocity 3D Image Analysis software (PerkinElmer) as previously reported^34^. Briefly, images were imported, timepoint sequenced, movement correct, background subtracted, and Gaussian filtered (fine). Regions of interest (ROIs) were generated based on the size of immunohistochemical stains using the active zone (AZ) marker brp (DSHB, nc82), and tiled (non-overlapping) over the area of GCaMP expression. Individual flashes, change in fluorescence (ΔF) 2 standard deviations above the background, were sorted into individual ROIs. ROI location (coordinates), and events were compiled and subsequently exported to excel for further analysis. Flashes were assigned to a single timepoint to prevent single events occurring over multiple frames from duplicate counts. The release probability for each ROI was calculated by dividing the total number of events detected within that ROI divided by the recording time (60 s). Release probability maps per ROI/AZ were generated in Microsoft Excel.

### Electrophysiology

Wandering third instar larvae were isolated from the side of fly vials, washed in HL3.0 saline. Larvae were dissected dorsal side up using previously established protocols^35^. Sharp intracellular recordings were conducted using 40-60 MΩ microelectrodes backfilled with 3M KCl. Recordings were amplified using an Axoclamp 2B amplifier (Molecular Devices) with a 0.1x headstage and digitized using a minidigi 1B (Molecular devices). Excitatory junctional potentials were elicited via a Master 8 stimulator (A.M.P.I) and a suction electrode (A-M Systems). For input resistance measurements, chord resistance was determined at the beginning and end of each trial, using 0.1, 0.2, 0.3, 0.4, and 0.5 nA. Signals were acquired using Axoclamp (11.2) and analyzed using Clampfit and Minianalysis. Data were exported to Microsoft Excel and figures were generated using GraphPad Prism.

### Muscle force recordings

Wandering third instar larvae isolated from the side of vials were dissected in HL3.0 saline. Larva were dissected dorsal side up, eviscerated, pinned down on the posterior end and the anterior end was attached to a fine minuten pin attached to the glass tube at the end of a 403A Force transducer system (Aurora Scientific). The brain was removed and the segmental nerve branches were sucked into a suction electrode (A-M systems). Nerve evoked contractions were generated using bursts of electrical stimuli from a Master 8 (A.M.P.I.) stimulator. Impulse duration was 5×10^−4^ seconds, interburst duration was kept constant at 15 seconds, burst durations were 600 milliseconds, and stimulation frequencies were stepped up from 1-to-150 Hz^36^. Each stimulation frequency was elicited 6 times and the corresponding voltage trace was acquired using Dynamic Muscle Acquisition software (Aurora Scientific). Digitized data were imported and processed in Matlab using custom code^36^.

### Analysis and Statistics

GraphPad Prism 11 was used for statistical analysis. Appropriate statistical metrics were performed for each dataset along with the F, and P statistics. Statistical comparisons were made with controls unless noted. Appropriate sample size was determined using a normality test. Data are presented as the mean ± SEM unless otherwise stated (\**p <* 0.05, \*\**p <* 0.01, \*\*\**p <* 0.001, n.s. = not significant).

## Results

### Expression of truncated pathogenic htt leads to intracellular aggregation in larval muscle fibers

To model htt expression in skeletal muscle, a pan muscle driver (Mef2-Gal4) was crossed with two transgenic *Drosophila* lines under the control of an upstream activator sequence (UAS). Two variants of the human htt protein were expressed with different glutamine repeats in the polyQ tract: a control (non-pathogenic, NP-htt) with 15 glutamine repeats and an experimental (pathogenic, P-htt) with 138 glutamine repeats using the pan muscle driver (Mef2-Gal4). Both constructs also contain an N-terminal red fluorescent protein (RFP) tag for visualization using fluorescence microscopy. Given that caspase-6 mediated cleavage of htt at amino acid position 588 has been demonstrated as a critical step leading to htt-aggregation in the nervous system (NS), we explored this truncation in the invertebrate equivalent of vertebrate skeletal muscle. The *Drosophila* larval neuromuscular junction (NMJ) is a widely used model for understanding the nervous system, and larval body wall muscle and adult flight muscle are incredibly powerful tools for investigations of muscle biology generally (**Fig. 1, A-B**)^32,36–38^. Anatomically, third instar *Drosophila* larvae are bilaterally symmetrical, and divided into 14 larval segments. Eight of the 14 segments comprise the bulk of the animal and are composed of repeated abdominal segments (**Fig. 1 B**). Each segment has 30 unique and individual muscle fibers, innervated by 31 motor neurons, all known and categorized (**Fig. 1 C-D**)^39^.

**Figure 1.**
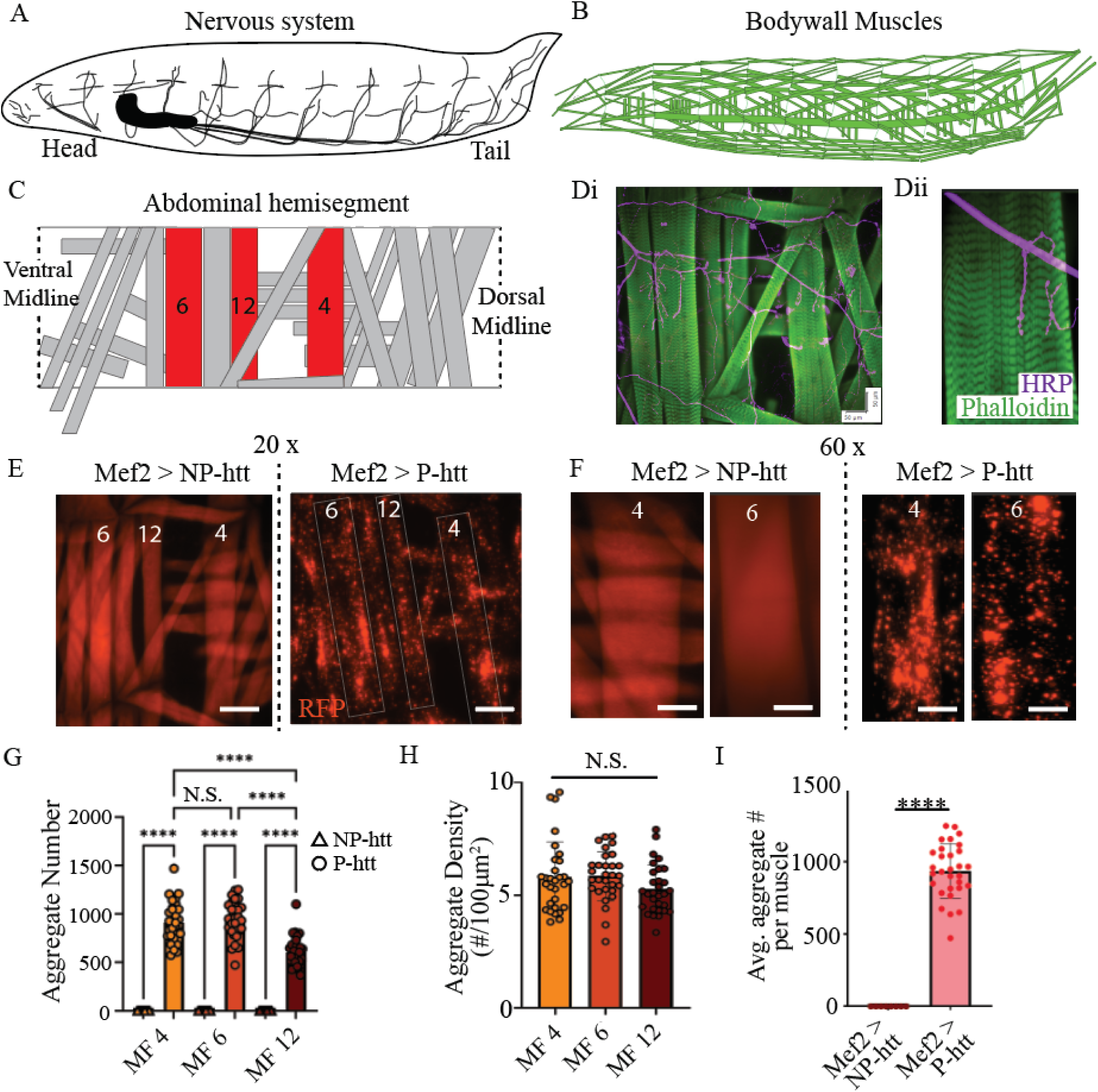
Pathogenic huntingtin expression in body wall muscle. Schematic of 3-dimnesional third instar larva highlighting the CNS in **A.** and body wall muscles in **B. C.** Model a single abdominal hemisegment in third instar larvae, each gray box illustrates the location and orientation of a different individual muscle fiber. The three individual body wall muscles, 4, 6, and 12, used through this study are highlighted in red. **D.** Fluorescence image of body wall muscles from a single abdominal hemisegment in a Canton S. larvae following immunohistochemistry against phalloidin (green) and horseradish peroxidase (HRP, magenta). Scale bars: Di: 50 microns Dii: 20 microns. **E.** Representative fluorescence images taken at 20x magnification from a single abdominal hemisegment highlighting muscle fibers 4, 6, and 12 acquired from Mef2-Gal4 > UAS-Q15-htt (Mef2 > NP-htt), and Mef2-Gal4 > UAS-Q138-htt (Mef2 > P-htt). Scale bar: 120 microns. **F.** Representative fluorescence images taken at 60x magnification from single abdominal muscle fibers 4, and 6 acquired from Mef2-Gal4 > UAS-Q15-htt (Mef2 > NP-htt), and Mef2-Gal4 > UAS-Q138-htt (Mef2 > P-htt). Scale bar: 50 microns. **G.** Total aggregate number per muscle from 3 different muscles, MF 4, MF 6, and MF 12, acquired from Mef2 > NP-htt and Mef2 > P-htt (N=30 per MF). **H.** Aggregate density (number of aggregates per 100 microns^2^, N=30 MFs) in Mef2 > P-htt for MF 4, 6, and 12. **I.** Average aggregate number per muscle in MF 4 from Mef2 > NP-htt and Mef2 > P-htt. ****P<0.0001.

Fluorescence imaging from an immunohistochemical stain from Mef2-Gal4 > UAS-htt-Q15 (Mef2 > NP-htt) highlights htt-protein accumulation in muscle fibers (MF) 4, 6, and 12 **(Fig. 1 E)**. In our previous investigation examining these htt constructs in the NS, we defined htt-aggregates as any non-motile fluorescent aggregate over 0.2 microns^2^^21^. In Mef2-Gal4 > UAS-htt-Q15 (Mef2 > NP-htt) no quantifiable RFP puncta were observed in 90 muscles imaged from 5 different animals (18 muscles per animal, 3 abdominal segments, MF 4, 6, and 12). However, in larvae from Mef2-Gal4 > UAS-htt-Q138 (Mef2 > P-htt) a dramatic amount of aggregation was observed in every muscle fiber examined (**Fig 1 E-F**). Aggregation was first quantified by tabulating the total number of aggregates observed from 30 muscle fibers from each of the 3 representative muscles (**Fig. 1 G**). We initially speculated upon a relationship between muscle size (volume) and total aggregate load, and found a significantly greater number of aggregates in MFs 4 and 6 compared to 12 (one-way ANOVA, F= 375.9, P=<0.0001, N=30, 5 animals per genotype). However, when we standardized for muscle area and calculated aggregate density per 100 micrometers^2^, no difference was observed across the 3 muscles analyzed (**Fig. 1 H**, one-way ANOVA, F= 2.0, P=0.14, N=30, 5 animals per genotype). Given the lack of statistical difference between the muscles with respect to aggregate density, our analysis focused on a single muscle fiber, MF 4, for the remainder of the study. MF 4 from Mef2 > NP-htt did not reveal any RFP-positive aggregates while Mef2 > P-htt showed significantly greater aggregate density (**Fig. 1 I**).

### Pathogenic htt aggregate morphology and fluorescence intensity are related to aggregate size

Next, we further characterized the size and distribution of aggregates from Mef2 > Q138 from MF 4. A total of 30 individual muscle fibers were analyzed (6 muscles fibers 4s from abdominal segments 3-5, from 5 different animals). A total of 27906 aggregates were found in these 30 muscle fibers, ranging in size from 0.2 to 61 *μ*m^2^. Aggregates were binned according to their size in increments of 1 *μ*m^2^ (**Fig 2. A**). More than half (56%) of the aggregates were less than 1 *μ*m^2^, and 93% of all aggregates were less than 5 *μ*m^2^. When examining the aggregates, a strong positive correlation between aggregate size and fluorescence intensity was noted (**Fig. 2** inset). Upon closer visual examination of individual aggregates of increasing size, the change in non-uniformity and increase in fluorescence is striking (**Fig 2 B-C).**

**Figure 2.**
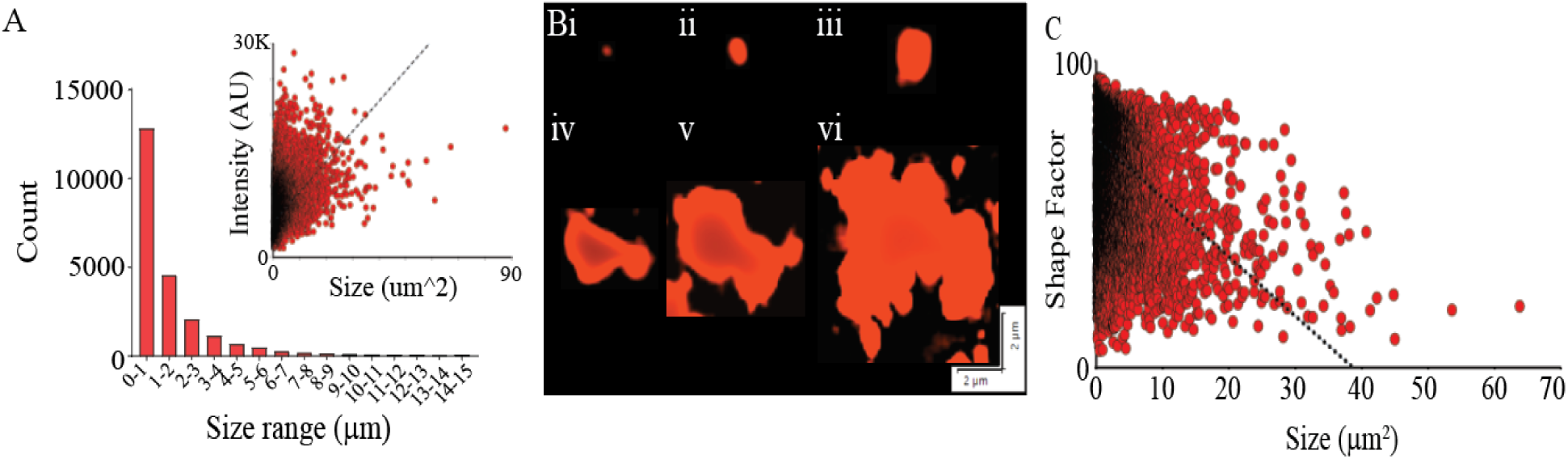
Htt aggregates increase in fluorescence intensity and become increasingly irregular in shape with increases in size. **A.** Compilation of all 27906 htt-positive aggregates from Mef2 > P-htt from MF 4 into bins increasing sequentially by 1 micron^2^ from left to right. Given their infrequency, aggregates over 15 microns^2^ were omitted. Inset: Aggregate size versu fluorescence intensity for all aggregates from Mef2 > P-htt. Line of fit: y=425x + 4939. **B**. Representative aggregates increasing in size from **i-v:** (size in microns & shape factor): **Bi:** 0.25 & 0.98, **Bii:** 4.88 & 0.81, **Biii:** 11.94 & 0.77, **Biv:** 17.35 & 0.66, **Bv:** 35.49 & 0.53, **Bvi:** 58.00 and 0.49. Scale bars: 2 microns. **C.** Aggregate size and shape factor for all 27906 htt-positive aggregates. Line of fit: y= -0.0193x + 0.75.

### Delaying pathogenic huntingtin expression leads to reduced intracellular accumulation

As HD is a progressive disorder, increasing in disease severity with time, we next explored the time-dependence of htt-aggregate accumulation in muscles. Using developmental time specific Gal4 driver lines we were able to induce htt expression at different stages of muscle development. Several different muscle driver lines have been identified which begin expressing at different stages of development. Mef2-Gal4 is reported to begin expressing at embryonic stage 12^40^ (other reports show expression as early as stage 7)^41^, while G7-Gal4 and C57-Gal4 are both reported to begin expressing during the 2^nd^ instar larval stage, with G7-Gal4 being a slightly stronger Gal4-driver (**Fig. 3 A**)^40^. 5053A-Gal4 is a Gal4-driver which expresses in a single abdominal muscle fiber within each hemisgment, MF 12, beginning at embryonic stage 8 (**Fig 3 A**)^42^. G7-Gal4 was crossed with NP-htt and P-htt to examine the developmental dependence of htt-aggregation in muscle. Similar to the Mef2 > NP-htt larval muscle, no RFP-positive aggregates were observed in third instar muscle from G7-Gal4 > UAS-htt-Q15 (**Fig. 3 Bi,** G7 > NP-htt). However, third instar larvae from G7-Gal4 > UAS-htt-Q138 (**Fig. 3 Bii-iii,** G7 > NP-htt) showed a substantial amount of RFP-positive aggregation in body-wall muscles. Compiling aggregates from 30 MF 4 from 5 animals generated a total of 5356 total aggregates, ranging in size from 0.2 to 43 microns^2^. Separating those 5356 aggregates into 1 micron^2^ bins revealed a similar distribution to that observed for Mef2 > P-htt, with 44% of all aggregates being less than 1 micron^2^ (**Fig 3 C**, Mef2>P-htt: 56%), and 89 % of aggregates less than 5 microns^2^ (Mef2>P-htt: 93%). While still a strikingly large number of RFP-positive puncta in muscles, G7>P-htt showed an 80% reduction in aggregate number compared to Mef2>P-htt (5356 vs. 27906). Figure 3 D shows the significant reduction in aggregate number per muscle for 30 MF 4 from Mef2 > P-htt compared to G7 > P-htt (t-test, P<0.0001, N=30 from 5 animals per genotype). Figure 3 E demonstrates a significant reduction in aggregate density for G7>P-htt compared to Mef2 > P-htt (t-test, P<0.0001, N=30 from 5 animals per genotype). Similar to what was observed from the aggregates from Mef2>P-htt, the aggregates appeared to become more irregularly shaped as the size of the aggregate increased (Fig. 3 F).

**Figure 3.**
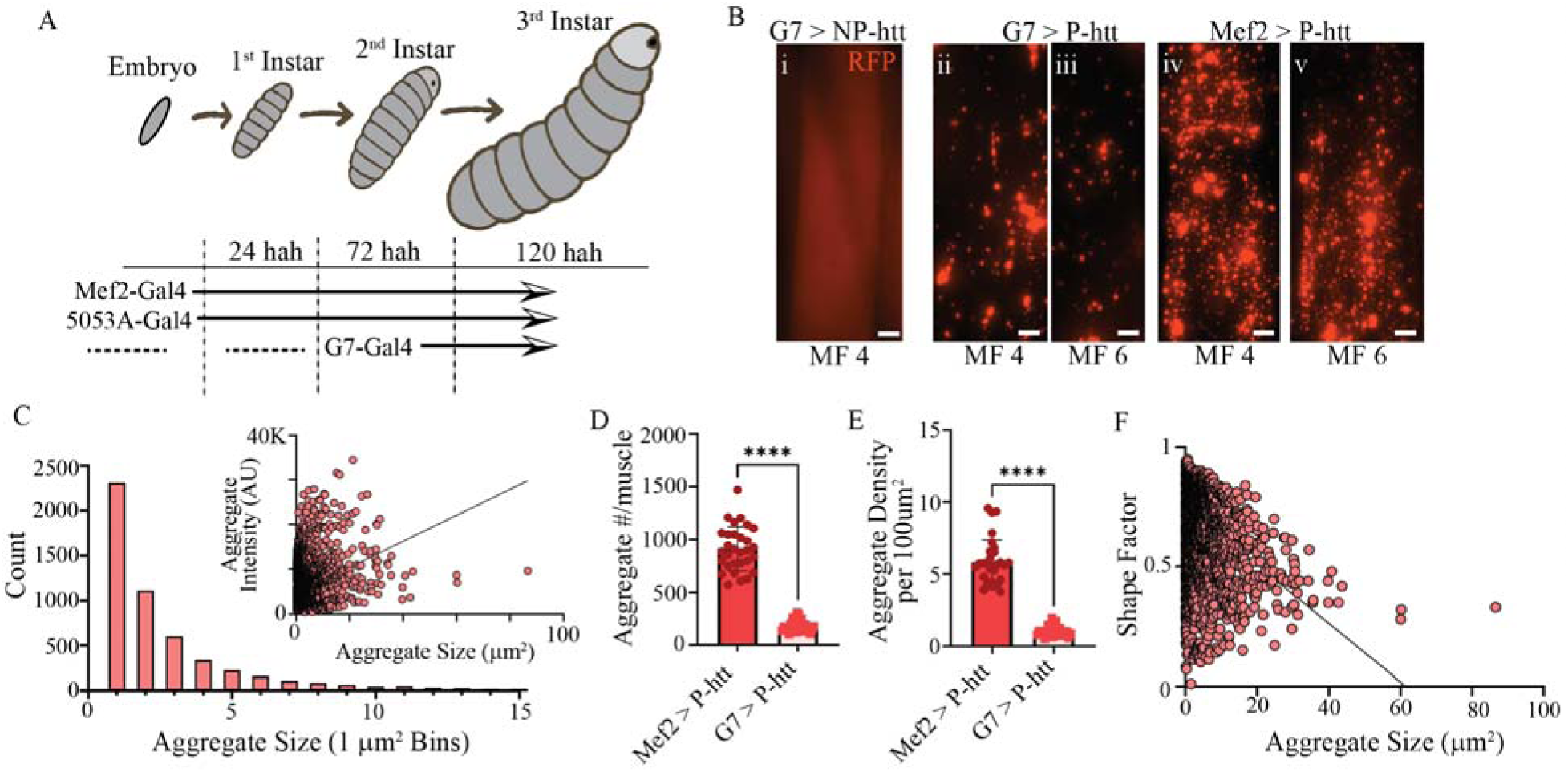
Developmental timing of htt-expression critically impacts aggregate formation. **A.** Schematic of developmental progression of *Drosophila* from embryo to third instar larvae, including the hours after hatching (hah) to reach each larval stage, and when each of the 3 different muscle driver lines begins expressing. **B.** Representative fluorescence microscopy images of **Bi:** MF 4 from G7-Gal4 > UAS-Q15-htt (G7 > NP-htt), **Bii-iii:** MFs 4 and 6 from G7-Gal4 > UAS-Q138-htt (G7 > P-htt) and **Biv-v:** Mef2-Gal4 > UAS-Q138-htt (Mef2 > P-htt). Scale bar: 20 microns. **C.** Aggregate size of 5336 RFP-positive puncta measured from 30 different MF 4 in G7 > P-htt. Aggregates were binned in 1-micron bins starting at 0-1, up-to 14-15. Aggregates greater than 15 were omitted given their infrequency. Inset: Aggregate size versu fluorescence intensity for each of the 5336 aggregates measured. **D.** A comparison of aggregate number per muscle for 30 individual MF 4s from Mef2 > P-htt and G7 > P-htt. **E.** Comparison of aggregate density per 100 micron^2^ from each of the 30 muscle fibers measure from Mef2 > P-htt and G7 > P-htt. ****P<0.0001. **F.** Aggregate size vs shape factor for 5336 RFP-positive aggregates measure from 30 different MF 4 in G7 > P-htt. Shape factor formula: 4π x area/perimeter^2^. ****P<0.0001.

Driving the expression of non-pathogenic, UAS-htt-Q15 with the single muscle driver 5053A-Gal4 (5053 > NP-htt) did not reveal any RFP-positive aggregation in MF 12 (Fig 3 G). Using 5053A-Gal4 to drive pathogenic, UAS-htt-Q138 (5053A>P-htt) did show aggregate accumulation in muscle fiber 12, but in no other muscles (e.x. MF 4 vs 12, Figure 3 H). RFP expression in MF 12 in 5053A>P-htt was dramatically reduced compared to MF 12 from Mef2>P-htt animals (SF 1). These data indicated that developmental timing is a critical determinant for aggregate number and size to increase.

### Gross muscle morphology and sarcomere structure were not altered by pathogenic htt expression

Given the profound aggregate density observed in Mef2>P-htt larvae, we next posited a probable impact on muscle structure and function. Phalloidin (α-F-actin) immunostaining was conducted on Canton S, Mef2>NP-htt, Mef2>P-htt, and G7>P-htt animals to examine changes in actin/sarcomere structure (**Fig 4A-C)**. Whole muscle fiber length and width measurements were taken from 30 MF 4, and the area of each MF 4 was computed. No changes in MF length, width, or area were observed across the 5 genotypes (**Fig. 4 D:** length: one-way ANOVA, F=2.5, P>0.05; width: one-way ANOVA, F=2.4, P>0.05, area: one-way ANOVA, F=2.7, P>0.05). Estimates of sarcomere and I-band lengths were determined from profile lines running anterior-posterior (**Fig. 4 C**). Similar to what was observed for whole muscle fibers, no change in sarcomere length (**Fig. 4 E:** one-way ANOVA, F= 12.11, P>0.05) and I-band length (**Fig 4 F:** one-way ANOVA, F=2.2, P>0.05) were observed. Immunostaining revealed a modification in nuclear shape within muscle fibers, which was particularly apparent upon Mef2 induced expression of P-htt. Those modifications in nuclear shape where not accompanied by any alterations in nuclei number per muscle fiber.

**Figure 4.**
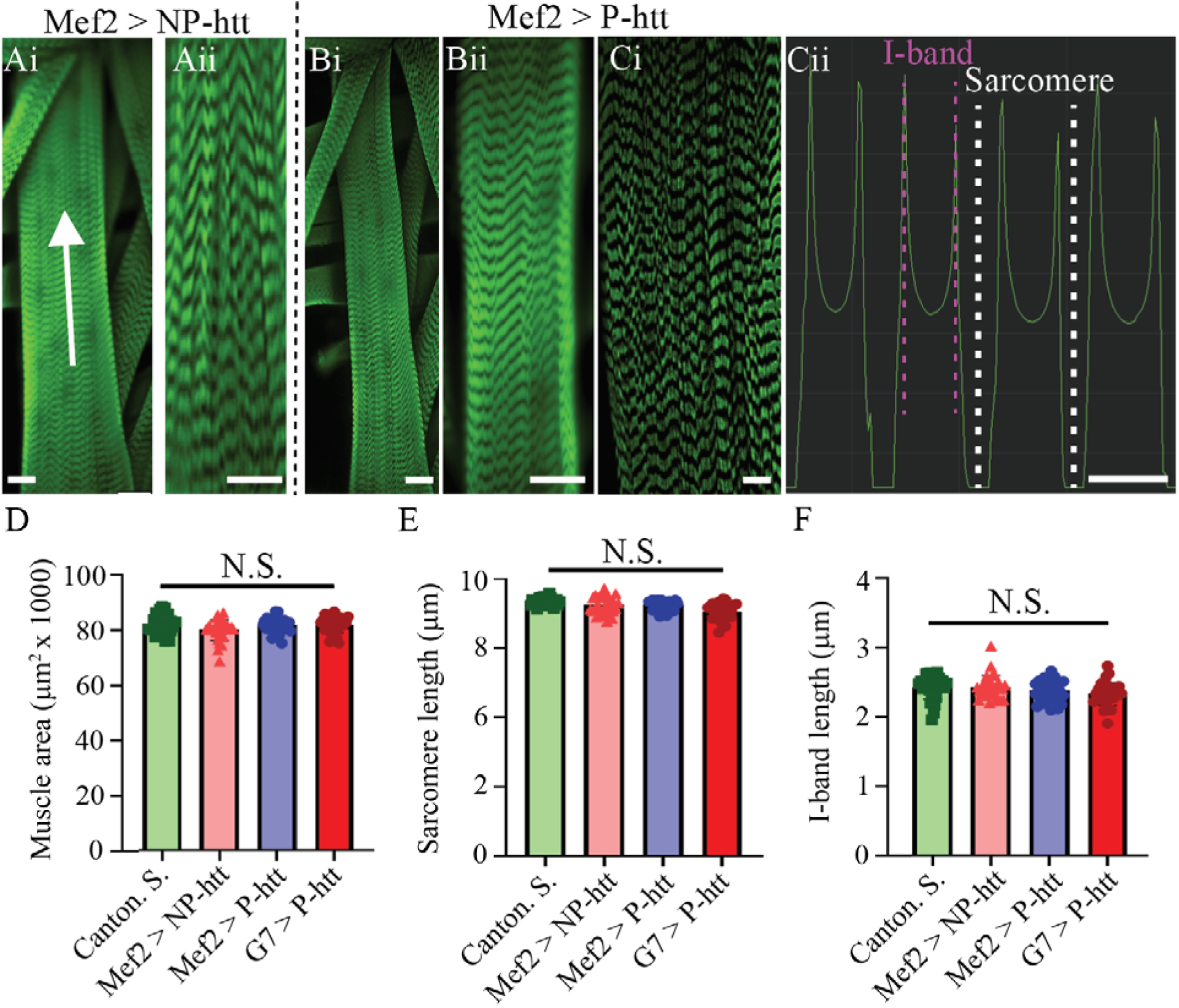
Pathogenic huntingtin expression does not change muscle macrostructure or myofilament structure. Representative immunostained images of muscle fiber 4 from **A:** Mef2 > NP-htt, and **B:** Mef2 > P-htt. **i:** 20x images of MF 4, **ii:** 60x images from MF4. **C:** Screen-capture from Olympus CellSens software using the line profile tool drawn from posterior-to-anterior of animal along MF 4, showing fluorescence intensity from a phalloidin-stained animal. Quantiifcations of sarcomere length and I-band length were made using this line profile tool. **D:** muscle area (muscle length times width) from 30 individual MF 4 from each of the four genotypes. Measurements of **E:** sarcomere length and **F:** I-band length from 30 different MF 4 from each of the four genotypes using the line profile tool.

### Myonuclear morphology is altered by pathogenic htt aggregation in a time-dependent manner

Next, a comprehensive assessment of muscle nuclei was conducted using DAPI immunostaining. Under low-magnification (20 x objective) the total number of nuclei per muscle was measured and no statistical differences were observed between the 5 genotypes (**Fig. 5 A**, One-way ANOVA, F=2.2, P>0.05, N=30 MF from 5 animals). However, the nuclei appeared misshapen, and dimmer in the Mef2>P-htt MF compared to Mef2>NP-htt. To increase precision and efficacy of nuclear shape analysis we imaged nuclei at higher resolution (63X, **Fig 5. B-C)**. At this magnification it became strikingly apparent the Mef2>P-htt nuclei were significantly larger in area compared to Mef2>NP-htt or Canton S. controls (**Fig. 5 D-F**, One-way ANOVA, F=156.6, P<0.0001). Interestingly G7>P-htt were also significantly greater in area compared to Mef2>NP-htt and Canton S, but significantly smaller than Mef2>P-htt (**Fig. 5 F**, One-way ANOVA, F=156.6, P<0.0001), suggestive of an intermediate phenotype. DAPI fluorescence can provide a good estimate of DNA content/density as it binds stoichiometrically to DNA^43^. Nuclei from MF 4 in Mef2>P-htt larvae showed a statistically reduced fluorescence intensity value compared to Mef2>NP-htt and Canton S. controls (**Fig. 5 G**, One-way ANOVA, F=711.2, P<0.0001). G7>P-htt again revealed an intermediate phenotype with respect to DAPI fluorescence intensity being statistically greater than G7>NP-htt and Canton S. controls, but statistically lesser than Mef2>P-htt (**Fig 5 G**). DAPI-stained nuclei from wild-type, Canton S. larvae have a symmetrical, round appearance, which was measured using the shape factor analysis in Olympus Cellsens software to be on average 0.9 (**Fig 5 D, H**). Nuclei from Mef2>P-htt larvae were typically irregularly shaped, generating a shape factor value significantly different from Mef2>NP-htt or Canton S (**Fig. 5 D, E, H**, One-way ANOVA, F=38.5, P<0.0001). Nuclei from G7>P-htt larval MFs did not show a significant change in shape factor compared to G7>NP-htt or Canton S. controls (**Fig 5 H**).

**Figure 5.**
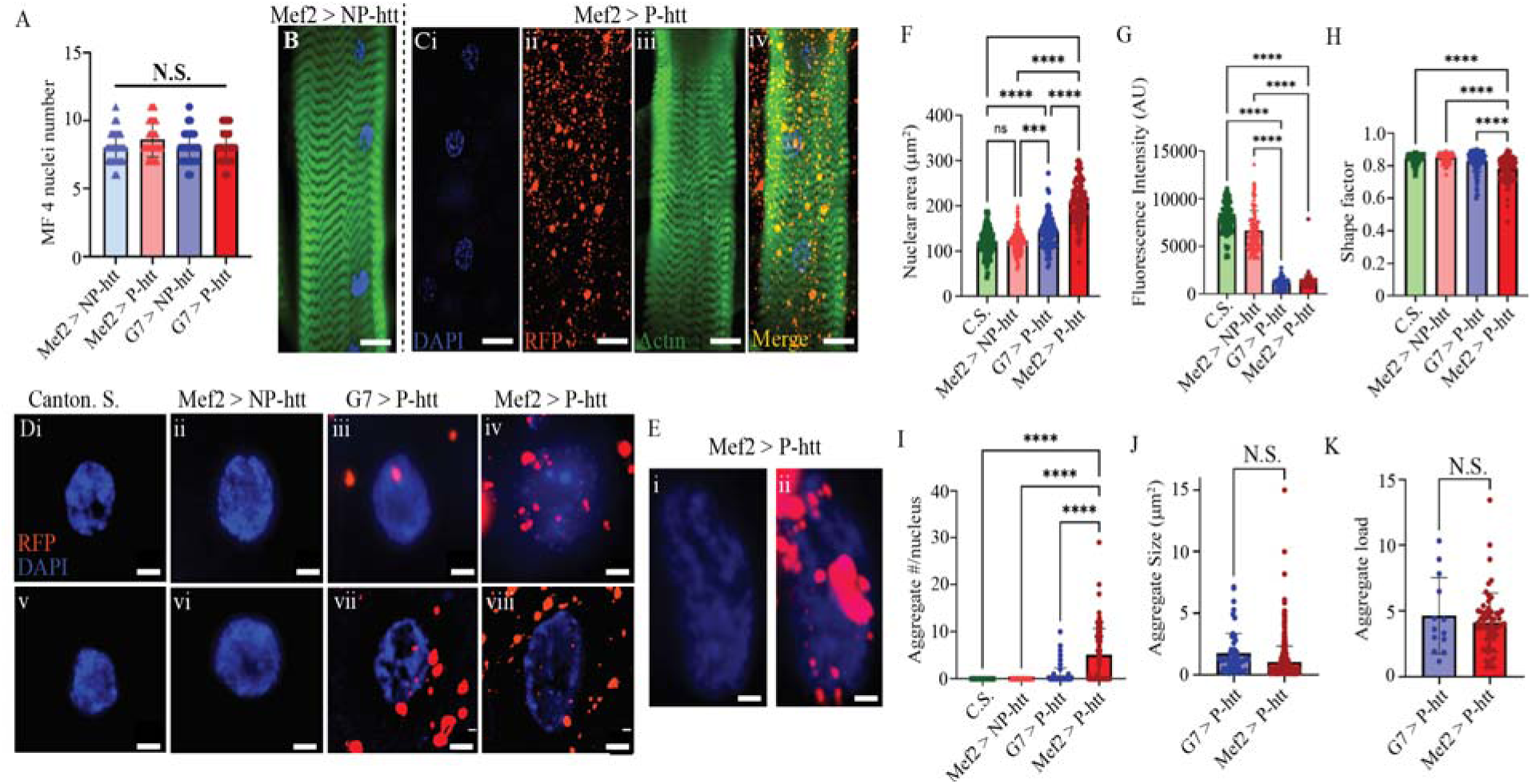
Pathogenic htt expression leads to htt-protein aggregates in the nucleus, and a dramatic change in nuclear shape. **A.** Over-expression of non-pathogen (NP-htt) or pathogenic huntingtin (P-htt) using either muscle driver G7-Gal4 or Mef2-Gal4 did not cause a change in nuclear number per muscle fiber. Immunostain of MF 4 from **B:** Mef2 >NP-htt, and **C:** Mef2 > P-htt. Green: phalloidin, blue, DAPI, red: RFP. **Ci-iv:** individual fluorescence channels are shown from **i-iii**, and a merge of all three channels in **iv.** Representative immunostained images from nuclei in MF 4 control (**Di, v**), Mef2 > NP-htt (**bii, vi**), G7 > P-htt (**Diii, vii**), and Mef2 > P-htt (**iv, vii**). Blue: DAPI, red: RFP. **E:** Representative nucleus from a Mef2 > P-htt showing a dramatic change in nuclear shape, intensity, and htt-protein aggregation (**Ei**: DAPI only, **Eii**: DAPI + RFP). Quantification of changes in: **F**: fluorescence intensity, **G:** shape factor, and **H:** nuclear area from each of the four genotypes. Nuclear RFP-positive htt-aggregates were quantified from each of the four genotypes. I: The total number of aggregates from each nucleus. J: The size of each aggregate from G7 > P-htt and Mef2 > P-htt. The total aggregate load percentage (combined area of htt-positive aggregates per nucleus divided by the nuclear area) of each aggregate from G7 > P-htt and Mef2 > P-htt. ***P<0.001, ****P<0.0001.

We next postulated that the change in nuclear shape/structure in muscle nuclei expressing P-htt may be due to htt-aggregate inclusions within the nuclei. Z-stacks of nuclei from MF 4 from all genotypes were imaged for DAPI and RFP, and overlayed revealing the presence of RFP-positive puncta within the nuclei only for those animals expressing P-htt (**Fig. 5 Div, Dvii, Eii**). Given the lack of RFP-positive aggregates in Canton. S. and Mef2>NP-htt, a statistical difference in intranuclear htt-aggregate number between Mef2-Gal4 and G7-Gal4 driving P-htt was observed (**Fig 5 I**, One-way ANOVA, F=, P<0.05). Next, intranuclear aggregate size from Mef2-Gal4 and G7-Gal4 driving P-htt was computed for all intranuclear aggregates, and no significant difference was observed (**Fig. 5 J,** t-test, df=65.3, P>0.05). Lastly, we examined whether a correlation was observed between nuclei size and aggregate load (total area of htt-aggregates divided by nuclear area x 100, **Fig. 5 K,** t-test, df=15.3, P>0.05). Here again no statistical difference was found, which may indicate an upper threshold or mechanism for intranuclear misfolded protein clearance. It is noteworthy that many of the nuclei from Mef2>P-htt appeared to have numerous and large areas devoid of DAPI immunostaining (**Fig. 5 D, E**). However, no obvious correlation existed between the presence of these dark areas and RFP-positive puncta.

### Pathogenic htt aggregates colocalize with mitochondrial accumulations in body wall muscles

Previous work from cultured skeletal muscle and dissociated muscle tissue from mice/human models of HD, have generated conflicting data whether metabolic changes occur in muscle fibers expressing P-htt^6,9,44^. To assess carbohydrate metabolism, we first conducted a periodic-acid Shifff (PAS). Stains from these animals did not reveal any obvious differences in carbohydrate metabolism between Mef2>P-htt, Mef2>NP-htt, and Canton S. larvae (data not shown). Next an Oil-Red-O stain was conducted to look for changes in lipid metabolism in animals expressing pathogenic huntingtin in larvae body wall muscle. Here again, no obvious differences in lipid staining were observed between Mef2>P-htt, Mef2>NP-htt, and Canton S. larvae (data not shown). Next, an immunohistochemical stain was conducted against the alpha subunit of mitochondrial ATP synthase (ATP5α), to examine changes in the location and structure of mitochondria within the larval muscle. Similar to previous reports in *Drosophila*, the ATP5α immunostain reveals faint banding patterns, and accumulation around nuclei (**Fig. 6 A**)^45^. However, in G7>P-htt and Mef2>P-htt the immunostaining was dramatically different as large and numerous GFP-positive puncta from the ATP5α immunostain were scattered across the muscle (**Fig. 6 B-C**). The total number of ATP5α-aggregates was quantified from each of the 3 genotypes revealing statistical significance between each (**Fig. 6 D,** One-way ANOVA, F=56.3, P<0.0001, N=30 MF from 5 animals). The average size of ATP5α-aggregates was significantly different for Mef2>P-htt and G7>P-htt compared to C.S. but not between the two P-htt expressing lines (**Fig. 6 E,** One-way ANOVA, F=114.3, P<0.0001, N=30 MF from 5 animals). Total mitochondrial load (total area of ATP5α-aggregates per muscle) for each genotype was significantly different between Mef2>P-htt and G7>P-htt, and C.S. (**Fig 6 F,** F=87.5, P<0.0001, N=30 MF from 5 animals). A Pearson’s correlation revealed a strong overlap between the RFP-aggregates and mitochondrial marker for both lines expressing P-htt (Mef2>P-htt: 0.65, G7>P-htt: 0.71). Collectively these data reveal a structural change in mitochondria in muscles expressing pathogenic human huntingtin.

**Figure 6.**
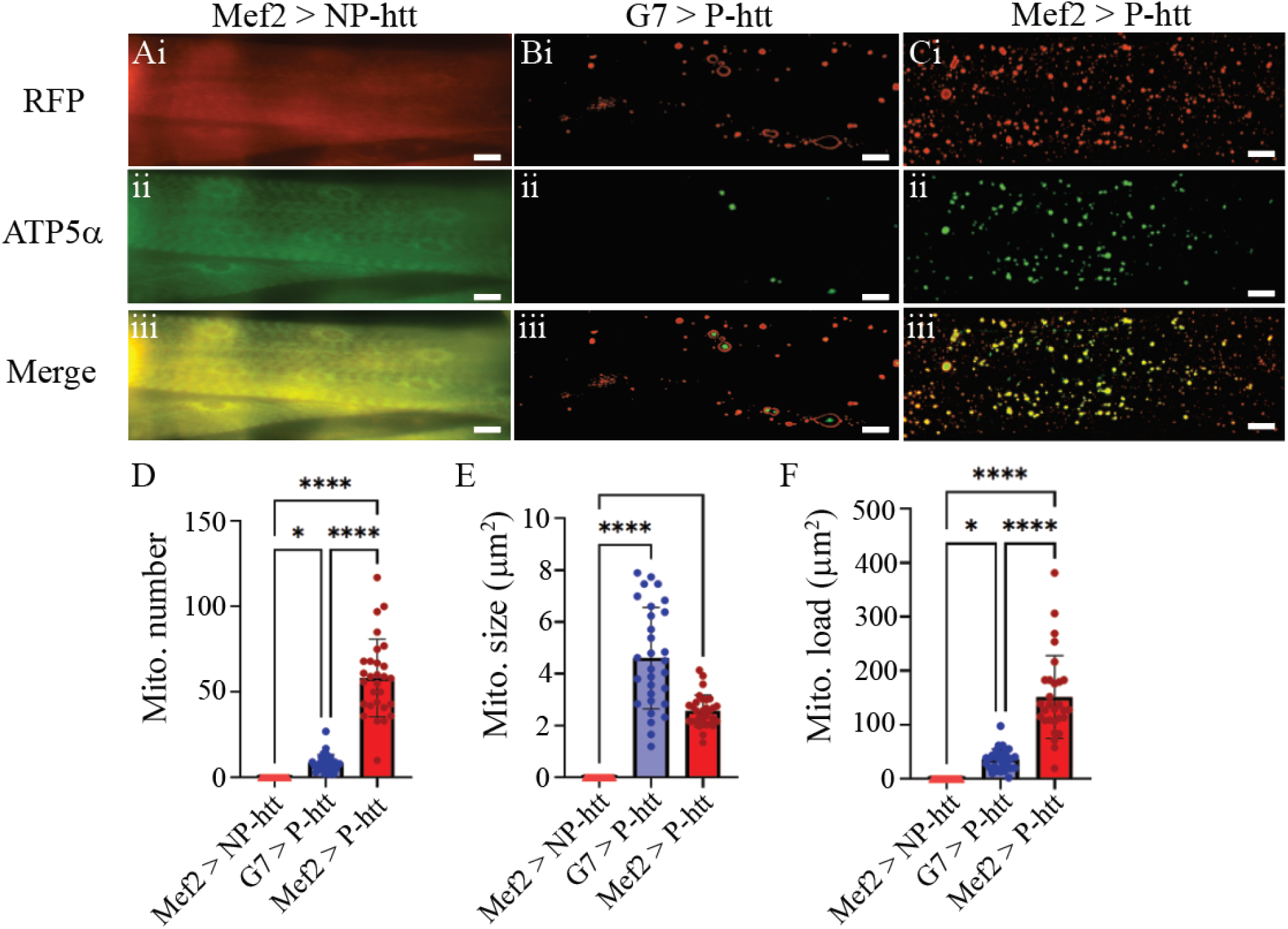
P-htt expression leads to mitochondrial aggregate formation in muscles. Representative immunostained images from columns **A**: Mef2 >NP-htt, **B**: G7 > P-htt, and **C**: Mef2 > P-htt. Row **i**: RFP-only fluorescence channel, **ii**: ATPα only channel, and **iii**: merge of both channels. Quantification of changes in ATP5α (mitochondria:mito) **D:** aggregate number, **E:** aggregate size and **F:** total mitochondrial aggregate load (combined ATP5α area). *P<0.05, ****P<0.0001.

### NMJ ultrastructure is not impacted by pathogenic huntingtin in muscles

During development a dynamic relationship exists between motor neurons and muscles, where motor neuron derived growth cones are guided towards an appropriate target along the surface of the muscle which secretes guidance cues, while muscles are extending myopodia to initiate a physical interaction between the two cells, leading to the formation of the NMJ, subsequently subject to synaptic pruning thereafter^38^. Given Mef2 expression begin during early embryonic development the possibility for disruptions in the formation of the NMJ are possible. Additionally, the NMJ is a dynamic structure with continuous molecular turnover, homeostatic changes, and physical stresses from larval movement^38,46,47^. Thus, the integrity of the NMJ connection was assessed via anti-HRP immunostaining (neuronal membrane marker, **Fig. 7 A**) and anti-dlg (postsynaptic scaffolding marker, **Fig. 7 B**). From the anti-hrp immunostain, the total innervation length along the surface of MF 4 was examined from Mef2>P-htt, Mef2>NP-htt, and Canton S. larvae and no significant differences were observed (**Fig 7 C**, One-way ANOVA, F=2.6, P>0.05, N=30 MF from 5 animals). The total number of boutons was also measured using the anti-hrp immunostain, and no significant differences were observed (One-way ANOVA, F=0.46, P=0.6, N=10 MF from 3 animals). The extent of postsynaptic density along the surface of MF 4 was also measured from the 3 genotypes and no significant difference was observed (**Fig. 7 D**, One-way ANOVA, F=3.1, P>0.05). One interesting observation was numerous dlg ‘islands’, where dlg-positive immunostaining of NMJs occurs independently from typical main innervation along the surface of MF 4 (**Fig. 7 Bii-iii**, top right). Taken together, no obvious change in pre- or postsynaptic NMJ structure has resulted from pathogenic human huntingtin expression in these larvae muscle.

**Figure 7.**
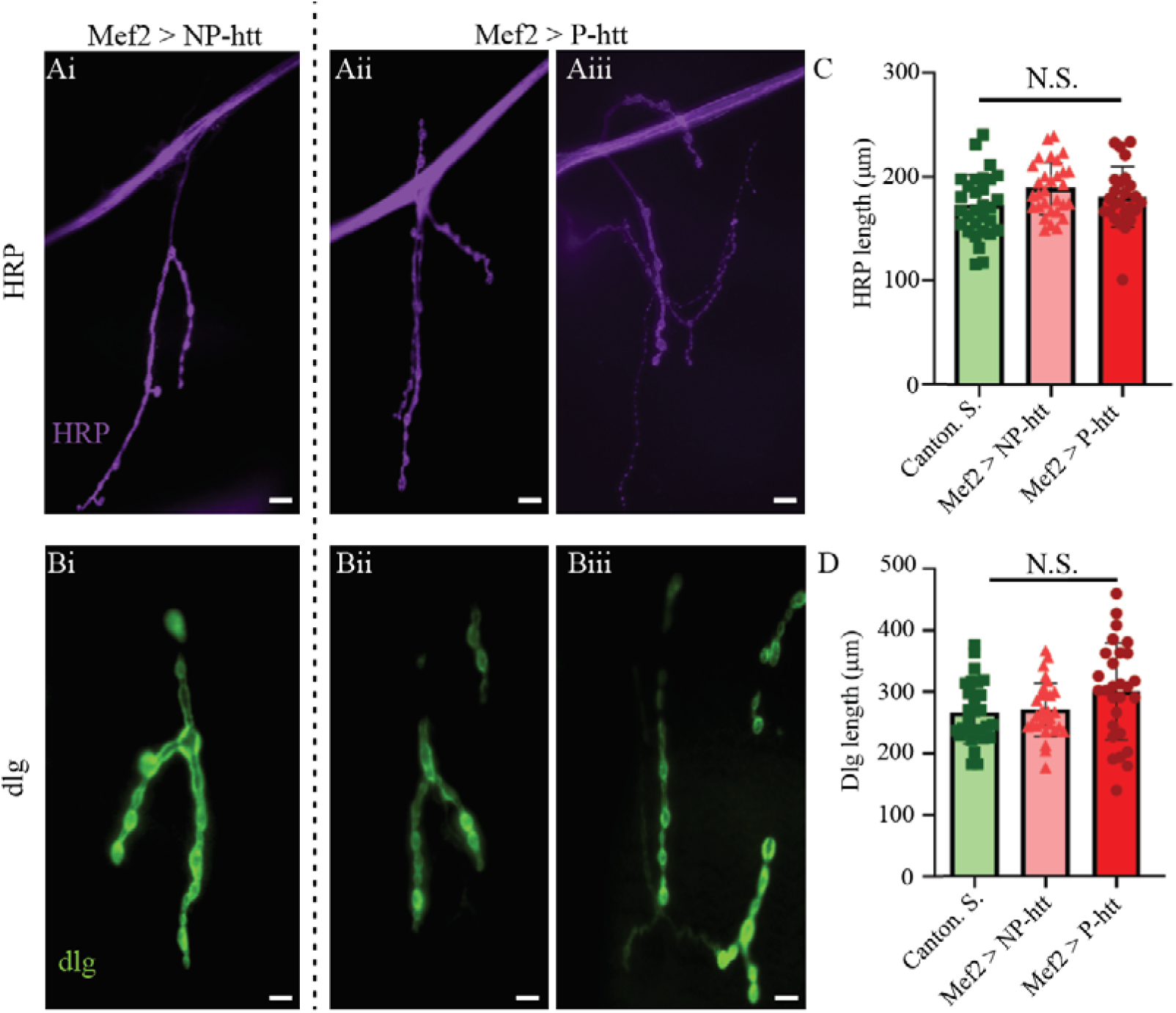
Pathogenic htt expression did not impact neuromuscular junction structure pre- or postsynaptically. Immunostaining of larval NMJs at MF 4 from Mef2>NP-htt (**Ai, Bi**) and Mef2>P-htt (**Aii-iii, Bii-iii**) with horseradish peroxidase (HRP, **A**) and discs-large (dlg, **B**). Quantification of length of HRP (**C**) and dlg (**D**) along the surface of MF 4 from Canton S., Mef2> NP-htt, and Mef2>P-htt.

### Muscle expression of mutant huntingtin causes synaptic transmission deficits

While no structural changes were observed at the NMJ, we next assessed whether pathogenic human huntingtin expression leads to functional changes in neuromuscular transduction. Frist, intracellular electrophysiological recordings were obtained from MF 4 from Mef2>P-htt, Mef2>NP-htt, and Canton S. larvae (**Fig. 8 A**). Representative raw traces of evoked excitatory junctional potential (EJP) activity can be seen in Figure 8 B where severed motor neurons were stimulated at low frequency (0.2 Hz). Muscle-specific expression of P-htt resulted in a significant reduction in EJP amplitude compared to C.S. and Mef2>NP-htt (One-way ANOVA, F=1.8, P<0.0001, N=5 animals). The kinetics of EJPs were also examined to look for changes in rise or decay time (*τ*) and no significant differences were observed (Fig. 8 Bii: rise *τ*, One-way ANOVA, F=, P>0.05; **Biii**: decay *τ*, One-way ANOVA, F=1.6, P>0.05). Similarly, the amplitude and frequency of miniature end plate potentials (mini’s) were not significantly different between the three genotypes (**Fig. 8 Bv:** mini frequency: One-way ANOVA, F=0.9, P>0.05; **Bvi:** mini amplitude: One-way ANOVA, F=3.2, P>0.05)

**Figure 8.**
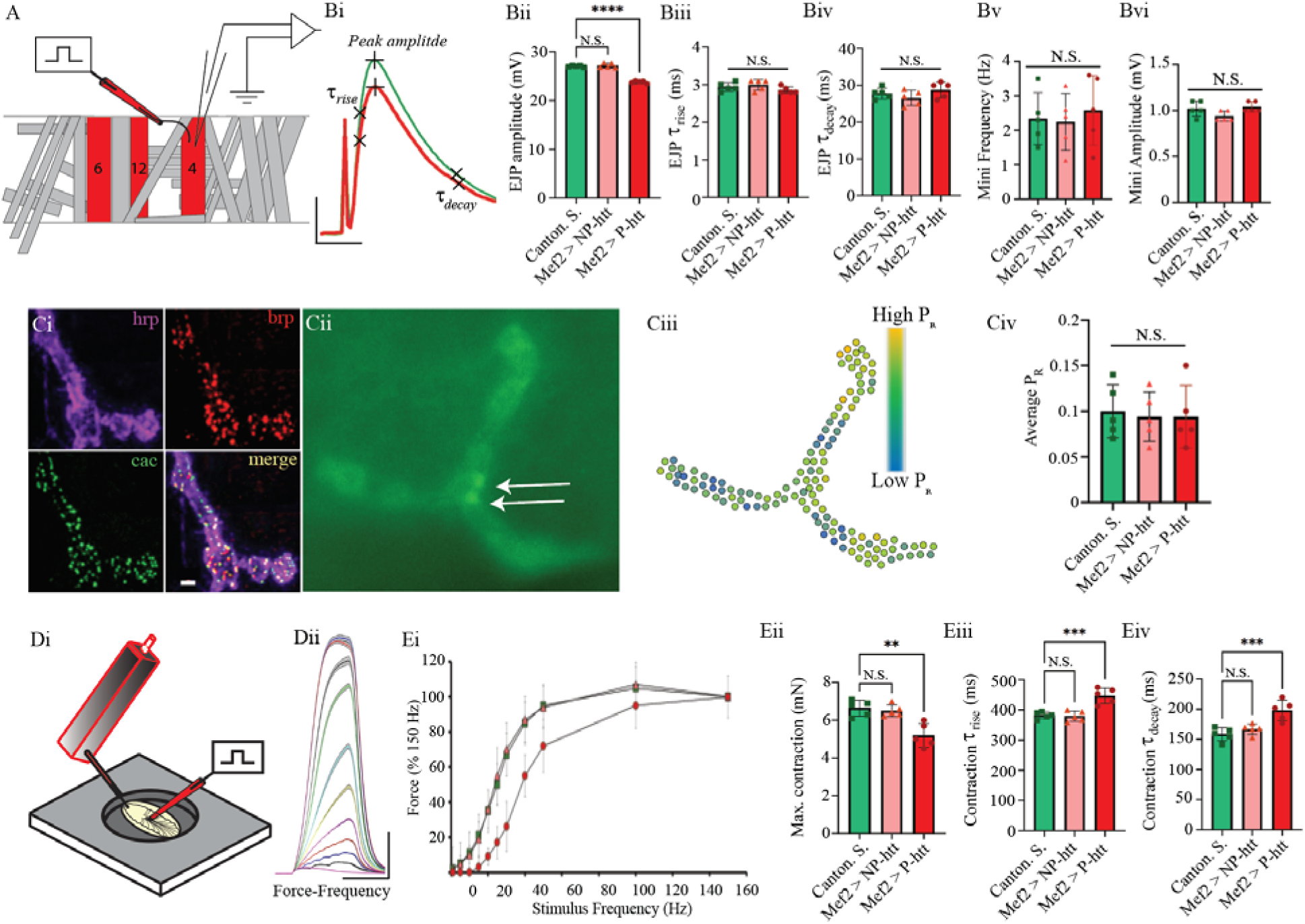
Pathogenic htt expression reduces neuromuscular transduction and excitation-contraction coupling. **A**. Schematic of third instar larval abdominal hemisegment highlighting muscle fibers 4, 6, 12 and location of suction electrode and intracellular microelectrode used for eliciting and recording EJPs. **Bi**. Representative electrophysiological EJP traces from Mef2>NP-htt (green) and Mef2>P-htt (red). Scale bars: 10 ms, 10 mV. Quantification of EJP amplitud (Bii), EJP rise tau (Biii) decay tau (Biv), mini frequency (**Bv**) and mini amplitude (**B**vi) from Canton S., Mef2>NP-htt, and Mef2>P-htt. **Ci.** Immunostaining of larval NMJ co-stained with hrp (magenta), cacophony (green), brp (red). **Cii**: Single frame from a Mef2> GCaMP, highlighting two calcium influx events (flashes, white arrows). **Ciii**. Release probability map from 1 min (600 frames) of a GCaMP recording (from Cii). Individual ROIs are color coded according to their release probability. **Civ:** Average release probability (P_R_) from all active zones over 1 min recording from all three genotypes. **Di**. 3D schematic of the experimental setup highlighting the force transducer setup. **Dii.** Six replicate contractions from each stimulation frequency (1, 5, 10, 15, 20, 25, 30, 40, 50, 100, 150) were averaged, overlayed, and plotted with 95% CI (grey). **Di.** Muscle force *vs*. stimulation frequency for the 3 genotypes. **Dii.** Representative muscle contraction from 40 Hz stimulation from Mef2>NP-htt (green) and Mef2>P-htt (red). Scale bars: 500 ms, 2 mN. Quantification of muscle contraction amplitude (**Diii),** rise tau (**Div**) and decay tau (**Dv**) from Canton S., Mef2>NP-htt, and Mef2>P-htt.

To examine possible changes at individual active zones, a myristoylated version of the genetically encoded calcium sensor (UAS-myr-GCaMP8s) was co-expressed in muscles with the non-pathogenic human huntingtin construct (Mef2-Gal4 > UAS-Q15-htt, UAS-myr-GCaMP8s) or with the pathogenic version (Mef2-Gal4 > UAS-Q138-htt, UAS-myr-GCaMP8s). Live fluorescence imaging was conducted from MF 4, and subsequently a release probability (P_r_) map was constructed from 1 minute of GCaMP activity at active zone (AZ) level resolution (**Fig. 8 C**). The P_r_ was averaged across all AZs per NMJ from 5 different animals per genotype, and no significant change in P_r_ was observed between the 3 genotypes (**Fig 8. Civ,** One-way ANOVA, F=0.1, P>0.05).

### Muscle force production and temporal dynamics are altered in larvae expressing P-htt

Lastly, excitation-contraction coupling was examined in larvae expressing P-htt using our novel force transducer with 10 micronewton resolution (**Fig. 8 Di**)^36^. Force-frequency curves were generated for each animal by stimulating motor neurons at step-wise increases in stimulation frequency (1, 2, 5, 10, 15, 20, 2, 30, 40, 50, 100, and 150 Hz) while keeping the burst constant at 600 ms (**Fig. 8 Dii**). From these recordings, force-frequency curves were generated to depict how force increases with stimulation frequency from 1-to-150 Hz (**Fig. 8 Ei**). Canton S. and Mef2>NP-htt generated stereotyped force-frequency curves, with relatively small contractions at low stimulation-frequencies which increase tremendously from 20-50 Hz, and saturate at and above 100 Hz. Mef2>P-htt animals failed to elicit muscle contraction below 15 Hz stimulation, which was followed by exponential increased in force with increases in frequency until plateauing at 100 Hz. In the exponential phase of the force-frequency curve, there was a clear deviation with lesser generation of force from 20-50 Hz compared to C.S. or Mef2>NP-htt animals. The maximal contraction elicited by larvae at 150 Hz stimulation was compared and found to be significantly different (One-way ANOVA, F=, P<0.0001). Interestingly, when individual contractions are overlayed, a noticeable change in the kinetics of contractions is observed (**Fig. 8 Eii**). A significant increase in both the rise (**Fig. 8 Eiv**, One-way ANOVA, F=20.3, P<0.001) and decay (**Fig. 8 Ev**, One-way ANOVA, F=13.5, P<0.001) time (*τ*) was observed. Thus, the effect of P-htt expression in muscles significantly reduced synaptic efficacy which directly translated to a comparable reduction in muscle contraction. However, the kinetics of muscle contractions were slowed in animals expressing P-htt compared to controls. Given the lack of change in sarcomeric structure, yet profound change in mitochondrial structure and distribution, the time in force kinetics are likely a result of calcium handing or ATP availability.

## Discussion

Huntingtin aggregation is a hallmark of HD pathology. In neurons, P-htt misfolds and forms oligomers and inclusion bodies that disrupt transcriptional regulation, proteostasis, mitochondrial function, and intracellular trafficking^1^. Similar aggregation phenomena have been observed in skeletal muscle from HD patients and animal models, suggesting shared mechanisms of proteotoxic stress across tissues^6^. However, muscle fibers are uniquely specialized cells characterized by high energetic demand, multinucleation, and tightly regulated protein turnover^48^. Skeletal muscle is a principal regulator of whole-body metabolism, accounting for the majority of insulin-stimulated glucose uptake and serving as a key site of oxidative phosphorylation and lipid utilization^49^. HD patients frequently exhibit hypermetabolism, unintended weight loss, insulin resistance, and impaired mitochondrial function which are metabolic disturbances that cannot be fully explained by central nervous system pathology alone^27^. Importantly, huntingtin is ubiquitously expressed, and many of the enzymes that aberrantly cleave htt in neurons such as caspases and calpains, are similarly expressed in peripheral tissues^50^. Therefore, it is possible that the physiological dysfunction resultant of pathogenic huntingtin accumulation could not only be recapitulated in peripheral tissue, but may also give rise to tissue-specific pathology not observed in the nervous system. Here we explored the impacts of P-htt expression in muscle cells an *in vivo* model organism.

The expression of pathogenic human huntingtin (P-htt) in *Drosophila* larval body wall muscle drives extensive intramuscular accumulation of huntingtin (htt) protein, consistent with the well-established aggregation-prone nature of expanded polyglutamine (polyQ) tracts^19,21,51,52^. This extensive aggregation is consistent with the well-established propensity of mutant huntingtin to misfold and assemble into higher-order inclusions, reflecting a breakdown in cellular proteostasis mechanisms^50,53–55^. Expression of P-htt in *Drosophila* larval body wall muscle provides a powerful *in vivo* model to examine tissue-specific proteostasis failure. In this context, P-htt accumulates as intracellular protein aggregates, reflecting its intrinsic propensity to misfold and form β-sheet–rich oligomers and inclusions^50,55^. The progressive buildup of these aggregates in muscle fibers suggests that the protein quality control systems, particularly the ubiquitin–proteasome system and autophagy, are either overwhelmed or functionally impaired^54^. Muscle tissue, being highly metabolically active and structurally dependent on cytoskeletal integrity, may be especially vulnerable to such disruptions^54,55^. The high aggregate burden suggests that protein quality control systems, including the ubiquitin–proteasome pathway and autophagy, are either insufficient or overwhelmed in this tissue context. *In vitro* analyses show proteosome dysfunction following even mild expansions in the polyQ length (∼40) but becoming increasingly dysfunctional particularly at longer polyQ repeat lengths (∼120+)^56–59^. Similarly, autophagosome trafficking dynamics and subsequent involvement in degradation are significantly impaired by pathogenic htt. Such accumulation is likely to impose both physical and biochemical stress on muscle cells, creating a toxic intracellular environment that disrupts normal cellular function. The relatively uniform aggregate density across different muscle fibers suggests that P-htt toxicity is largely cell-autonomous and not restricted to specific subsets of myofibers^60^. The broad size distribution of aggregates, from ∼0.2 to >60 μm², with a predominance of smaller species (<5 μm²), is consistent with a dynamic aggregation process involving nucleation, growth, and coalescence^1^. Our observation that larger aggregates exhibit increased fluorescence intensity supports progressive recruitment of soluble or oligomeric P-htt into inclusions, while their increasingly irregular morphology suggests structural heterogeneity and possible coalescence of smaller aggregates into less ordered assemblies^61–63^. Additionally, htt aggregates have been demonstrated to act as indiscriminate sinks, trapping various organelles, which could further contribute to the irregular morphologies as aggregate size increases. Such changes in shape and packing may reflect biophysical transitions between aggregate states, which have been proposed to differentially influence toxicity^64^.

Despite this substantial aggregate burden, the preservation of sarcomere organization, as well as unchanged horseradish peroxidase (HRP) and discs-large (Dlg) immunostaining, indicates that core cytoskeletal architecture and neuromuscular junction (NMJ) structural integrity remain intact^20,38,65^. Furthermore, the absence of detectable changes in periodic acid-Schiff staining and Oil Red O staining suggests that glycogen and neutral lipid stores are not grossly perturbed, arguing against widespread metabolic storage defects at this stage^66,67^. Together, these findings indicate that P-htt aggregation does not immediately compromise global muscle architecture or energy storage, but instead exerts more selective effects on specific intracellular systems. This dissociation between aggregate accumulation and gross structural disruption is consistent with prior work suggesting that early pathogenic events in HD models often spare overt cellular architecture while impairing more subtle aspects of cellular function^68,69^. The lack of detectable changes in these markers further implies that P-htt toxicity in muscle is not mediated by immediate breakdown of contractile or synaptic scaffolding, but instead reflects selective vulnerability of specific intracellular systems. In contrast, mitochondrial organization is profoundly altered, with mitochondrial aggregates showing nearly complete spatial overlap with P-htt inclusions. This striking co-localization is consistent with extensive evidence linking mutant huntingtin to mitochondrial dysfunction, including impaired trafficking, altered fission–fusion dynamics, and deficits in bioenergetics^70,71^. The sequestration or clustering of mitochondria within P-htt-rich regions may disrupt their normal distribution along muscle fibers, limiting efficient ATP delivery and calcium buffering at sites of high demand^72^. Moreover, physical association between aggregates and mitochondria may impair mitochondrial quality control pathways, such as mitophagy, thereby exacerbating oxidative stress, energetic failure, and protein misfolding repair/removal pathways^72–75^. These findings support the idea that mitochondrial dysregulation is a central and early feature of P-htt -toxicity, even in non-neuronal tissues. Despite growing recognition of peripheral pathology in HD, the mechanistic relationship between pathogenic huntingtin aggregation and metabolic dysfunction in skeletal muscle remains incompletely defined. It is unclear whether aggregation directly drives bioenergetic impairment, whether metabolic stress promotes aggregation, or whether these processes reinforce one another. Addressing this gap is essential for understanding the full pathophysiological spectrum of HD and for designing therapeutic strategies that target both neuronal and peripheral tissues.

Transcriptional dysregulation of metabolic regulators, such as PGC-1α and downstream oxidative phosphorylation genes, have been reported in HD muscle, potentially coupling proteotoxic aggregation to bioenergetic failure^76^. Impairments in autophagy and ubiquitin, proteasome pathways may further exacerbate aggregate accumulation, creating a feed-forward cycle of metabolic stress and defective protein clearance. Given the central role of skeletal muscle in systemic energy homeostasis, such muscle-intrinsic defects may contribute significantly to the hypercatabolic state observed in HD. Nuclear abnormalities further underscore the multi-compartmental impact of P-htt expression. Enlarged, irregularly shaped nuclei with reduced DAPI fluorescence intensity are indicative of altered chromatin organization and compromised nuclear integrity^77^. Such phenotypes are consistent with previous reports demonstrating that mutant huntingtin interacts with transcriptional regulators and chromatin-modifying complexes, leading to widespread transcriptional dysregulation^78,79^. Reduced DAPI intensity may reflect changes in chromatin compaction or accessibility, while nuclear deformation suggests defects in nuclear lamina structure or nucleo-cytoskeletal coupling^80,81^. The emergence of irregularly shaped nuclei points to compromised structural stability of the nuclear envelope and cytoskeletal coupling. P-htt aggregates localized to or near the nucleus may mechanically distort nuclear shape or interfere with nucleo-cytoskeletal linkages, such as the LINC complex^82,83^. Such deformation is often associated with impaired gene regulation, altered mechanotransduction, and increased susceptibility to DNA damage^81^. Moreover, the polyQ fibrils appear to physically pierce the nuclear envelope. The ruptures created do not fully repair as the P-htt fibrils remain embedded in the nuclear membrane^53^. Together, these nuclear phenotypes support a model in which pathogenic huntingtin exerts toxicity not only through cytoplasmic aggregation but also by directly perturbing nuclear organization, structure, and function, ultimately contributing to muscle cell dysfunction. Collectively, these changes point to a disruption of nuclear homeostasis that may further amplify cellular dysfunction through altered gene expression programs in cells expressing P-htt.

At the functional level, P-htt expression leads to a significant reduction in excitatory junctional potential (EJP) amplitude without changes in EJP kinetics, indicating a selective impairment in synaptic strength rather than timing. This phenotype is consistent with reduced neurotransmitter release, impair vesicle cycling or decreased postsynaptic responsiveness, while preserved kinetics suggest that receptor gating and ion channel dynamics remain largely intact^84,85^. The maintenance of normal temporal properties alongside reduced amplitude highlights a deficit in the efficacy of neuromuscular transmission rather than its coordination. Importantly, these synaptic deficits are accompanied by a marked reduction in muscle force production and a slowing of both contraction and relaxation kinetics, as reflected by increased rise-and decay-*τ* values^36^. Resultantly, synaptic deficits are correlated or possibly compounded by intrinsic impairments in contractile function. P-htt-expressing muscles exhibit significantly reduced force production along with prolonged force rise and decay kinetics^36^. These changes point to defects in excitation–contraction coupling, potentially involving disrupted calcium handling, impaired sarcoplasmic reticulum function, or reduced efficiency of cross-bridge cycling. Previous studies indicate that Store-Operated Calcium Entry (SOCE) acts as a critical mechanism for replenishing sarcoplasmic reticulum calcium stores, and in skeletal muscle, this process is heavily reliant on the structural and functional interaction between STIM1, Orai1, and junctophilin^86,87^. The convergence of mitochondrial dysfunction, nuclear abnormalities, and weakened synaptic input provides a multifaceted explanation for the observed decline in muscle performance. Collectively, these findings underscore the complex, multi-compartmental nature of P-htt toxicity in muscle tissue, where selective subcellular disruptions precede and likely drive functional impairment.

Collectively, these data support a model in which pathogenic huntingtin exerts selective yet multifaceted effects in muscle cells. These findings also highlight the cell-autonomous toxicity of P-htt outside of neuronal systems, reinforcing the idea that Huntington’s disease is a multi-system disorder. Aggregate formation in larval muscle may sequester essential proteins, disrupt nuclear architecture, or alter gene expression through aberrant interactions with transcriptional regulators. Additionally, the dynamic nature of aggregate growth suggests a balance between nucleation, expansion, and potential clearance mechanisms that shifts over time toward accumulation. Widespread protein aggregation occurs in the absence of gross structural disruption, but is tightly linked to mitochondrial clustering and nuclear abnormalities, ultimately leading to deficits in synaptic efficacy and contractile performance. This convergence of proteostatic failure, organelle dysfunction, and impaired neuromuscular function aligns with broader models of Huntington’s disease pathogenesis, in which early subcellular perturbations precede overt degeneration^88^. Recent medical advances have significantly improved the ability to target mutant huntingtin protein within the central nervous system, with technologies like siRNA and antisense oligonucleotides showing promise in lowering toxic protein levels. However, the current study and many others demonstrate HD is a multi-system disorder, not just a brain disease. Pathogenic huntingtin is expressed ubiquitously, and its accumulation in peripheral tissues contributes significantly to the profound symptoms and overall pathology. The *Drosophila* larval muscle system thus provides a valuable platform for dissecting how aggregate properties, such as size, density, and morphology, relate to cellular dysfunction, and for identifying modifiers that may restore proteostasis and preserve muscle function. The multifaceted impact of P-htt in muscle tissue highlights the importance of targeting intracellular pathways in all affected cells and tissues to mitigate disease-related dysfunction.

## Acknowledgements

Research supported by the National Institute of General Medical Sciences of the National Institutes of Health under award number R15GM155985. Generous financial support to KGO from the University of Toledo. Thank you to Dr. Anthony Scibelli for assistance in MATLAB code for muscle force analyses, and Dr. Yulia Akbergenova for assistance with GCaMP analysis and manuscript discussions. Thank Troy Littleton for generously providing UAS-htt-Q138 and UAS-htt-Q15 fly lines.

## Supplementary figures

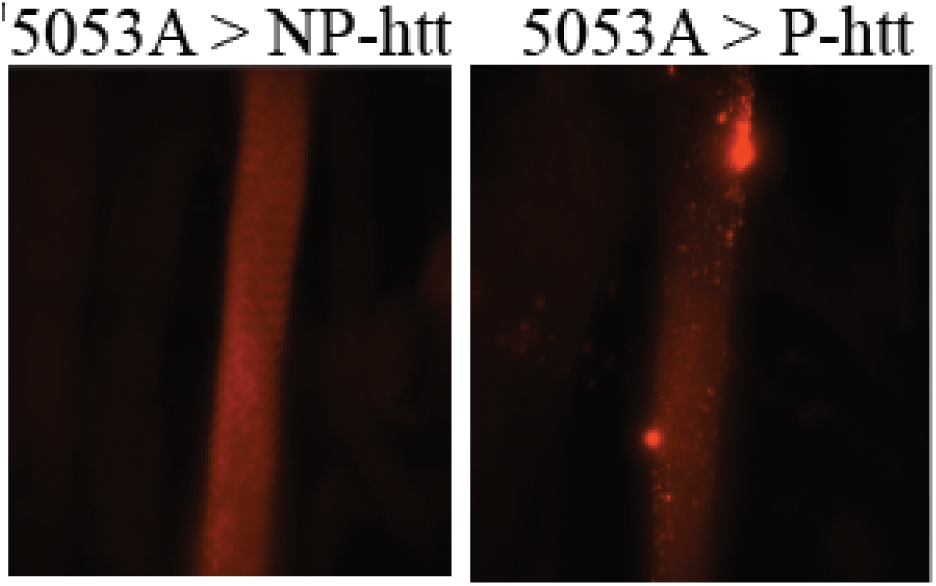
Representative fluorescence microscopy images of MF12 from 5053A-Gal4 > UAS-Q15-htt (5053A > NP-htt) and 5053A-Gal4 > UAS-Q138-htt (5053A > P-htt).

## References

1. Arrasate, M. & Finkbeiner, S. Protein aggregates in Huntington’s disease. Exp. Neurol. 238, 1–11 (2012).

2. Ross, C. A. et al. Huntington disease: natural history, biomarkers and prospects for therapeutics. Nat. Rev. Neurol. 10, 204–216 (2014).

3. Rüb, U. et al. Huntington’s disease (HD): the neuropathology of a multisystem neurodegenerative disorder of the human brain. Brain Pathol. 26, 726–740 (2016).

4. Mielcarek, M. Huntington’s disease is a multi-system disorder. Rare Diseases 3, e1058464 (2015).

5. Ogilvie, A. C., Nopoulos, P. C. & Schultz, J. L. Quantifying the Onset of Unintended Weight Loss in Huntington’s Disease: A Retrospective Analysis of Enroll-HD. J. Huntingtons Dis. 10, 485 (2021).

6. Bozzi, M. & Sciandra, F. Molecular Mechanisms Underlying Muscle Wasting in Huntington’s Disease. Int. J. Mol. Sci. 21, 8314 (2020).

7. Chang, C. P., Wu, C. W. & Chern, Y. Metabolic dysregulation in Huntington’s disease: Neuronal and glial perspectives. Neurobiol. Dis. 201, (2024).

8. Gil-Salcedo, A., Lunven, M., Jacquemot, C., Massart, R. & Bachoud-Levi, A. C. Specific contribution of cognitive and motor impairments with functional capacity and dependence in Huntington’s disease. J. Neurol. 272, 224 (2025).

9. Zielonka, D., Piotrowska, I., Marcinkowski, J. T. & Mielcarek, M. Skeletal muscle pathology in Huntington’s disease. Front. Physiol. 5, 380 (2014).

10. Mielcarek, M. et al. HDAC4-Myogenin Axis As an Important Marker of HD-Related Skeletal Muscle Atrophy. PLoS Genet. 11, e1005021 (2015).

11. Romer, S. H. et al. A mouse model of Huntington’s disease shows altered ultrastructure of transverse tubules in skeletal muscle fibers. J. Gen. Physiol. 153, (2021).

12. Turner, C., Cooper, J. M. & Schapira, A. H. V. Clinical correlates of mitochondrial function in Huntington’s disease muscle. Mov. Disord. 22, 1715–1721 (2007).

13. Chaturvedi, R. K. et al. Impaired PGC-1α function in muscle in Huntington’s disease. Hum. Mol. Genet. 18, 3048 (2009).

14. De Aragão, B. C. et al. Changes in structure and function of diaphragm neuromuscular junctions from BACHD mouse model for Huntington’s disease. Neurochem. Int. 93, 64–72 (2016).

15. Bradford, J. W., Li, S. & Li, X. J. Polyglutamine toxicity in non-neuronal cells. Cell Res. 20, 400 (2010).

16. Mielcarek, M., Smolenski, R. T. & Isalan, M. Transcriptional Signature of an Altered Purine Metabolism in the Skeletal Muscle of a Huntington’s Disease Mouse Model. Front. Physiol. 8, 127 (2017).

17. Coffey, S. R. et al. Peripheral huntingtin silencing does not ameliorate central signs of disease in the B6.HttQ111/+ mouse model of Huntington’s disease. PLoS One 12, e0175968 (2017).

18. Weiss, K. R. & Littleton, J. T. Characterization of axonal transport defects in Drosophila Huntingtin mutants. J. Neurogenet. 30, 212–221 (2016).

19. Weiss, K. R., Kimura, Y., Lee, W. C. M. & Littleton, J. T. Huntingtin aggregation kinetics and their pathological role in a Drosophila Huntington’s disease model. Genetics 190, 581–600 (2012).

20. Akbergenova, Y., Cunningham, K. L., Zhang, Y. V., Weiss, S. & Littleton, J. T. Characterization of developmental and molecular factors underlying release heterogeneity at drosophila synapses. Elife 7, (2018).

21. Hana, T. A. et al. Developmental and physiological impacts of pathogenic human huntingtin protein in the nervous system. Neurobiol. Dis. 203, 106732 (2024).

22. Landles, C. & Bates, G. P. Huntingtin and the molecular pathogenesis of Huntington’s disease. EMBO Rep. 5, 958 (2004).

23. Joshi, D. C. et al. The role of mitochondrial dysfunction in Huntington’s disease: Implications for therapeutic targeting. Biomedicine and Pharmacotherapy 183, (2025).

24. Neueder, A. et al. Huntington’s disease affects mitochondrial network dynamics predisposing to pathogenic mitochondrial DNA mutations. Brain 147, 2009–2022 (2024).

25. Chivet, M. et al. Huntingtin regulates calcium fluxes in skeletal muscle. J. Gen. Physiol. 155, e202213103 (2022).

26. van der Burg, J. M., Björkqvist, M. & Brundin, P. Beyond the brain: widespread pathology in Huntington’s disease. Lancet Neurol. 8, 765–774 (2009).

27. Nambron, R. et al. A Metabolic Study of Huntington’s Disease. PLoS One 11, e0146480 (2016).

28. Noordermeer, J., Klingensmith, J., Perrimon, N. & Nusse, R. dishevelled and armadillo act in the wingless signalling pathway in Drosophila. Nature 367, 80–83 (1994).

29. Ranganayakulu, G. et al. A Series of Mutations in the D-MEF2 Transcription Factor Reveal Multiple Functions in Larval and Adult Myogenesis in Drosophila. Dev. Biol. 171, 169–181 (1995).

30. Zhang, Y. Q. et al. Drosophila Fragile X-Related Gene Regulates the MAP1B Homolog Futsch to Control Synaptic Structure and Function. Cell 107, 591–603 (2001).

31. Akbergenova, Y., Cunningham, K. L., Zhang, Y. V., Weiss, S. & Littleton, J. T. Characterization of developmental and molecular factors underlying release heterogeneity at Drosophila synapses. Elife 7, (2018).

32. Michael, A. H., Hana, T. A., Mousa, V. G. & Ormerod, K. G. Muscle-fiber specific genetic manipulation of Drosophila sallimus severely impacts neuromuscular development, morphology, and physiology. Front. Physiol. 15, (2024).

33. Banerjee, S. et al. The Drosophila melanogaster Neprilysin Nepl15 is involved in lipid and carbohydrate storage. Scientific Reports 2021 11:1 11, 2099- (2021).

34. Melom, J. E., Akbergenova, Y., Gavornik, J. P. & Littleton, J. T. Spontaneous and Evoked Release Are Independently Regulated at Individual Active Zones. The Journal of Neuroscience 33, 17253 (2013).

35. Ormerod, K. G., Krans, J. L. & Mercier, A. J. Cell-selective modulation of the Drosophila neuromuscular system by a neuropeptide. J. Neurophysiol. 113, 1631–1643 (2015).

36. Ormerod, K. G., Scibelli, A. E. & Littleton, J. T. Regulation of excitation-contraction coupling at the Drosophila neuromuscular junction. J. Physiol. 600, 349–372 (2022).

37. Ormerod, K. G., Jung, J. & Joffre Mercier, A. Modulation of neuromuscular synapses and contraction in Drosophila 3rd instar larvae. J. Neurogenet. 32, 183–194 (2018).

38. Aponte-Santiago, N. A., Ormerod, K. G., Akbergenova, Y. & Littleton, J. T. Synaptic Plasticity Induced by Differential Manipulation of Tonic and Phasic Motoneurons in Drosophila. J. Neurosci. 40, 6270–6288 (2020).

39. Hoang, B. & Chiba, A. Single-cell analysis of Drosophila larval neuromuscular synapses. Dev. Biol. 229, 55–70 (2001).

40. Zhao, Z. & Geisbrecht, E. R. Stage-specific modulation of Drosophila gene expression with muscle GAL4 promoters. Fly (Austin). 19, 2447617 (2025).

41. Ranganayakulu, G., Elliott, D. A., Harvey, R. P. & Olson, E. N. Divergent roles for NK-2 class homeobox genes in cardiogenesis in flies and mice. Development 125, 3037–3048 (1998).

42. Klapper, R. The longitudinal visceral musculature of Drosophila melanogaster persists through metamorphosis. Mech. Dev. 95, 47–54 (2000).

43. Gomes, C. J., Harman, M. W., Centuori, S. M., Wolgemuth, C. W. & Martinez, J. D. Measuring DNA content in live cells by fluorescence microscopy. Cell Div. 13, 6 (2018).

44. Dickson, E., Soylu-Kucharz, R., Petersén, Å. & Björkqvist, M. Hypothalamic expression of huntingtin causes distinct metabolic changes in Huntington’s disease mice. Mol. Metab. 57, 101439 (2022).

45. Wang, Z. H., Clark, C. & Geisbrecht, E. R. Analysis of mitochondrial structure and function in the Drosophila larval musculature. Mitochondrion 26, 33–42 (2016).

46. Goel, P. & Dickman, D. Distinct homeostatic modulations stabilize reduced postsynaptic receptivity in response to presynaptic DLK signaling. Nat. Commun. 9, (2018).

47. Rozenfeld, E., Ehmann, N., Manoim, J. E., Kittel, R. J. & Parnas, M. Homeostatic synaptic plasticity rescues neural coding reliability. Nat. Commun. 14, (2023).

48. Peron, S., Zordan, M. A., Magnabosco, A., Reggiani, C. & Megighian, A. From action potential to contraction: neural control and excitation-contraction coupling in larval muscles of Drosophila. Comp. Biochem. Physiol. A Mol. Integr. Physiol. 154, 173–183 (2009).

49. Merz, K. E. & Thurmond, D. C. Role of Skeletal Muscle in Insulin Resistance and Glucose Uptake. Compr. Physiol. 10, 785 (2020).

50. Schulte, J. & Littleton, J. T. The biological function of the Huntingtin protein and its relevance to Huntington’s Disease pathology. Curr. Trends Neurol. 5, 65 (2011).

51. Ross, C. A. & Poirier, M. A. Protein aggregation and neurodegenerative disease. Nat. Med. 10 **Suppl**, S10 (2004).

52. Saudou, F. & Humbert, S. The Biology of Huntingtin. Neuron 89, 910–926 (2016).

53. Korsten, G. et al. Nuclear poly-glutamine aggregates rupture the nuclear envelope and hinder its repair. J. Cell Biol. 223, (2024).

54. Singh, A., Phogat, J., Yadav, A. & Dabur, R. The dependency of autophagy and ubiquitin proteasome system during skeletal muscle atrophy. Biophys. Rev. 13, 203 (2021).

55. Hu, C., Lin, M., Wang, C. & Zhang, S. Current Understanding of Protein Aggregation in Neurodegenerative Diseases. Int. J. Mol. Sci. 26, 10568 (2025).

56. Wong, Y. C. & Holzbaur, E. L. F. The Regulation of Autophagosome Dynamics by Huntingtin and HAP1 Is Disrupted by Expression of Mutant Huntingtin, Leading to Defective Cargo Degradation. The Journal of Neuroscience 34, 1293 (2014).

57. Holmberg, C. I., Staniszewski, K. E., Mensah, K. N., Matouschek, A. & Morimoto, R. I. Inefficient degradation of truncated polyglutamine proteins by the proteasome. EMBO J. 23, 4307 (2004).

58. Raspe, M. et al. Mimicking proteasomal release of polyglutamine peptides initiates aggregation and toxicity. J. Cell Sci. 122, 3262–3271 (2009).

59. Holmberg, C. I., Staniszewski, K. E., Mensah, K. N., Matouschek, A. & Morimoto, R. I. Inefficient degradation of truncated polyglutamine proteins by the proteasome. EMBO J. 23, 4307 (2004).

60. Marchioretti, C. et al. Skeletal Muscle Pathogenesis in Polyglutamine Diseases. Cells 11, 2105 (2022).

61. Aktar, F. et al. The huntingtin inclusion is a dynamic phase-separated compartment. Life Sci. Alliance 2, e201900489 (2019).

62. Weiss, K. R., Kimura, Y., Lee, W. C. M. & Littleton, J. T. Huntingtin Aggregation Kinetics and Their Pathological Role in a Drosophila Huntington’s Disease Model. Genetics 190, 581 (2012).

63. Waelter, S. et al. Accumulation of Mutant Huntingtin Fragments in Aggresome-like Inclusion Bodies as a Result of Insufficient Protein Degradation. 10.1091/mbc.12.5.1393 12, 1393–1407 (2017).

64. Olzscha, H. et al. Amyloid-like aggregates sequester numerous metastable proteins with essential cellular functions. Cell 144, 67–78 (2011).

65. Harris, K. P., Zhang, Y. V., Piccioli, Z. D., Perrimon, N. & Troy Littleton, J. The postsynaptic t-SNARE Syntaxin 4 controls traffic of Neuroligin 1 and Synaptotagmin 4 to regulate retrograde signaling. Elife 5, (2016).

66. Prats, C. et al. An Optimized Histochemical Method to Assess Skeletal Muscle Glycogen and Lipid Stores Reveals Two Metabolically Distinct Populations of Type I Muscle Fibers. PLoS One 8, e77774 (2013).

67. Kanungo, S., Wells, K., Tribett, T. & El-Gharbawy, A. Glycogen metabolism and glycogen storage disorders. Ann. Transl. Med. 6, 474 (2018).

68. Scherzinger, E. et al. Huntingtin-encoded polyglutamine expansions form amyloid-like protein aggregates in vitro and in vivo. Cell 90, 549–558 (1997).

69. Li, S. H. & Li, X. J. Huntingtin-protein interactions and the pathogenesis of Huntington’s disease. Trends in Genetics 20, 146–154 (2004).

70. Shirendeb, U. P. et al. Mutant huntingtin’s interaction with mitochondrial protein Drp1 impairs mitochondrial biogenesis and causes defective axonal transport and synaptic degeneration in Huntington’s disease. Hum. Mol. Genet. 21, 406–420 (2012).

71. Reddy, P. H. & Shirendeb, U. P. Mutant huntingtin, abnormal mitochondrial dynamics, defective axonal transport of mitochondria, and selective synaptic degeneration in Huntington’s disease. Biochim. Biophys. Acta Mol. Basis Dis. 1822, 101–110 (2012).

72. Dong, H. & Tsai, S. Y. Mitochondrial Properties in Skeletal Muscle Fiber. Cells 12, 2183 (2023).

73. Li, W. et al. Molecular mechanisms of mitochondrial quality control. Transl. Neurodegener. 14, 45 (2025).

74. Zhou, Q. Y. et al. The crosstalk between mitochondrial quality control and metal-dependent cell death. Cell Death & Disease 2024 15:4 15, 299- (2024).

75. Banarase, T. A. et al. Mitophagy regulation in aging and neurodegenerative disease. Biophys. Rev. 15, 239 (2023).

76. McMeekin, L. J., Fox, S. N., Boas, S. M. & Cowell, R. M. Dysregulation of PGC-1α-Dependent Transcriptional Programs in Neurological and Developmental Disorders: Therapeutic Challenges and Opportunities. Cells 10, 352 (2021).

77. Singh, I. & Lele, T. P. Nuclear morphological abnormalities in cancer – a search for unifying mechanisms. Results Probl. Cell Differ. 70, 443 (2022).

78. Zuccato, C., Valenza, M. & Cattaneo, E. Molecular mechanisms and potential therapeutical targets in Huntington’s disease. Physiol. Rev. 90, 905–981 (2010).

79. Sugars, K. L. & Rubinsztein, D. C. Transcriptional abnormalities in Huntington disease. Trends in Genetics 19, 233–238 (2003).

80. Stephens, A. D. et al. Chromatin histone modifications and rigidity affect nuclear morphology independent of lamins. Mol. Biol. Cell 29, 220 (2018).

81. Kalukula, Y., Stephens, A. D., Lammerding, J. & Gabriele, S. Mechanics and functional consequences of nuclear deformations. Nat. Rev. Mol. Cell Biol. 23, 583 (2022).

82. Grima, J. C. et al. Mutant Huntingtin Disrupts the Nuclear Pore Complex. Neuron 94, 93–107.e6 (2017).

83. Jun, M. et al. Subcellular Force Imbalance in Actin Bundles Induces Nuclear Repositioning and Durotaxis. ACS Appl. Mater. Interfaces 15, 43387–43402 (2023).

84. Macleod, G. T. & Zinsmaier, K. E. Synaptic Homeostasis on the Fast Track. Neuron 52, 569–571 (2006).

85. Frank, C. A., Kennedy, M. J., Goold, C. P. P., Marek, K. W. & Davis, G. W. W. Mechanisms Underlying the Rapid Induction and Sustained Expression of Synaptic Homeostasis. Neuron 52, 663 (2006).

86. Michelucci, A., García-Castañeda, M., Boncompagni, S. & Dirksen, R. T. Role of STIM1/ORAI1-mediated Store-Operated Ca2+ Entry in Skeletal Muscle Physiology and Disease. Cell Calcium 76, 101 (2018).

87. Kiviluoto, S. et al. STIM1 as a key regulator for Ca2+ homeostasis in skeletal-muscle development and function. Skelet. Muscle 1, 16 (2011).

88. Ross, C. A. & Tabrizi, S. J. Huntington’s disease: From molecular pathogenesis to clinical treatment. Lancet Neurol. 10, 83–98 (2011).

